# Comprehensive studies of Head Maralla, Punjab, Pakistan vegetation for ethnopharmacological and ethnobotanical uses and their elaboration through quantitative indices

**DOI:** 10.1101/2020.11.16.384420

**Authors:** Muhammad Sajjad Iqbal, Muhammad Azhar Ali, Muhammad Akbar, Syed Atiq Hussain, Noshia Arshad, Saba Munir, Hajra Masood, Samina Zafar, Tahira Ahmad, Nazra Shaheen, Rizwana Mashooq, Hifsa Sajjad, Munaza Zahoor, Faiza Bashir, Khizra Shahbaz, Hamna Arshad, Noor Fatima, Faiza Nasir, Ayesha Javed Hashmi, Sofia Chaudhary, Ahmad Waqas, Muhammad Islam

## Abstract

Head Maralla is a non-recognized wetland with diversified flora that becomes focus of current studies. Whole plant or their parts are being used for treating various maladies and they are the integral part of livelihood in the area. Unique species including *Osmunda regalis* is used for blood and renal diseases purifier. Wild plant resources are key to traditional ethnobotanical knowledge based practices and industrial applications. Current study reports Head Marala inhabitant’s interaction with these resources and identify priorities at species and habitat level for conservation. Four sites viz., River Tavi, Upstream Chenab, River Manawarwala Tavi and Bhalolpur were designated to record traditional knowledge through questionnaire and interviews during field trips. One hundred nineteen (119) plant species were identified belonging to 54 families, of which 87 species were of dicot, 12monocots, 05ferns, and 4 species of bryophytes. Fifty percent of the plant species were utilized as a whole for therapeutic purposes, followed by leaves which had more than 20% usage of total consumption. Ailments viz., urination (14%) followed by cough (8%), cold (7%), stomach (6%), asthma (6%), constipation (5%), laxative (5%), diarrhea (4%) etc., were associated with vegetation. Bronchial disorders, pneumonia, dyspepsia, anthelmintic and kidney stones, etc., were also among other diseases commonly cured by traditional knowledge. Fifteen percent of vegetation contributes as fodder species consumed by local community for livestock while almost 17% of local plants were utilized for industrial purposes like timber, fuel, furniture, wooden pots and sports goods. In conclusion the ecosystem of Head Maralla is a complex of aquatic, terrestrial and agricultural land that is located on climatic and geographical divides, which further add to botanical interest as included many wetland habitats with unique diversity of plants. It is suggested to devise comprehensive conservation strategies to safe indigenous knowledge in systematic way for comprehending ecological services.

## Introduction

Plants are in use like food, medicine, fruit, vegetables, fodder, fuel and many more and continuously adding new things to the daily life. Their importance gains more and more attention as technological development opened new avenues to cater maximum benefits, likewise gums, resins, fibers, adhesives, stabilizer, softeners, emulsifiers, vitamins, oils, lubricants etc. Novel ingredients proved effective and safe in use. Among plants diversity prominently 300 species are used worldwide in food, pharmaceutical, cosmetic and perfume industry as natural pigments, flavoring agents and in traditional health care system [1]. Pakistan has unique recognition as having wide range of plants due to its climate and topography. This is also evident from the natural heritage and remnants collected from old civilizations; Indus, Texila-Gandhara and Mohenjo-Daro. Accordingly, sustainable development of TRM in developing countries requires serious investment and to meet health millennium goals by 2025 on medicinal plants conservation and traditional medicines along with local practitioners in PHC for provision of better healthcare [2].

Plant product development needed ethnobotanical assessment through conservation and protection of natural plant resources, therefore it plays important role in various world societies [1]. Twenty species out of 20,000 provides 90% of our food requirements. Consequently, global heritage is required to be preserved for future generations, as it provides guarantee for socioeconomic activities of an area by conserving local indigenous and traditional knowledge across the regions [3–5]. The uses of plants and their products especially herbal medicines are not only evident from developing countries but also becoming fashionable in richer countries in Europe and North America, where a market sector is growing at the pace of 10-20% annually.

Active ingredients from medicinal plants are being tested and used in western countries too. A simple illustration of synthetic drug versus plant extract is from the common foxglove (*Digitalis purpurea* L.) as derived for heart diseases ‘digoxin and digitoxin’. Similarly, the valuable pain killers, morphine and codeine are extracted from opium poppy (*Papaver somniferum* L.) [5]. Proper reports on ethnobotanical knowledge heritage have been documented since 1996 to onward in Pakistan [6–10]. Literature supported that Himalayan ranges housed 70% of plants and animals in wild, which fulfill 70-80% population traditional medicines health care [11–12]. Likely Head Maralla is also situated at foot hills of Himalaya housed as hub of many wild species. Ethnobotanical knowledge is a treasure that is not only restricted to rural areas, benefits are also evident from urban communities due to safer, cheaper and minimum side effects [13]. During an expedition, a Pansar (local medicinal plants’ store keeper) told that his daily sale is 120 dollars, at minimum level in the Hafizabad District. Another example is a tonic ‘humdard guti’ is very famous for digestion in newborns till the age of two years.

According to NIH, 400 plant species are being used extensively in traditional medicines. The Tibbi Pharmacopoeia of Pakistan has listed around 900 single drugs and about 500 compound preparations. 27 local herbal companies are registered for commercial production and their turnover is comparable to multinational in Pakistan. Unregistered Hakim Khanas are in hundreds spread throughout the country. Traditional healers are around 50,000 in numbers, including homeopaths. Herbal medication is known as The Unani System of Medicine in Pakistan. Hakims are practitioner awarded four years degree from Tibbia colleges that are >30. The Unani Drug Act is implemented by Government of Pakistan through Tibbi Council under the Ministry of Health. To document traditional knowledge from local inhabitants’ various reports have been published. Likewise, 26 species of vascular plants of Mianwali district (south western parts of the Punjab province) have been reported for medicine, furniture, agriculture implements and as food [3]. In another study, 49 species of 29 families from district Attock, Punjab were documented, utilized for medicinal purposes, according to the interviews by 10 Hakims and 80 people [4].

Moreover, high diversity of plants is existed in northern and northwestern parts of Pakistan [14]. 400-600 medicinal plants out of 5700 are estimated to exist in Pakistan [15]. Herbs like isphaghol, sweet fennel, black cumin, black seeds, bishops’ weed, milk thistle, Indian hemp, *Datura, Sassurea* and *Trianthma* sp., are extensively used locally. Whole or different parts of plant are usually used like, rhizome, fruits, leaves, root and bark to treat cough, asthma, cold, pneumonia, hepatitis, kidney stones, anti-malaria, small pox, heart and sexual disorders etc., therefore plants contain enormous wealth for a healthy life style. In this remote area, locals are depending on indigenous plant resources to supply a range of goods and services, including grazing for livestock and medicinal supplies. Due to remote location, rough terrain, topography and varied climatic patterns, many areas within these districts still remain unexplored for species distribution and various ecosystem services. Probably, most important role of indigenous plants is to achieve conservation of natural habitats. This is how related to the major contribution that they make livelihood in terms of health support, financial income, cultural identity and livelihood security. So, under Convention on Biological diversity, the fair and equitable sharing of benefits from bioprospecting and save guarding of intellectual property rights are required to be exercised [16].

Head Maralla, is floristically quite rich tropical regions of Punjab. Ethnobotanical study of this area has never been conducted. Climate of the area is subjected to extreme variation. Farmer community is dominating by profession whereas government employees stand second, young ones with low rate of literacy are interested to serve in foreign countries to enjoy high living standards. Women are housewives and a small number are serving in government. Both men and women are engaged (self-employed) in collection and trade of wild plants. Mostly plants are used for food, shelter and therapeutic agents while majority of the material goes useless. Post harvest processing based techniques are lacking behind like, scientific knowledge about the useable parts, cultural practices, management, harvesting, time of collection, processing and packaging lead to post harvest losses on one hand while reduction in national revenue on the other hand. Therefore, the indigenous knowledge is going to be depleted, hence ethnobotanical survey was planned and documented for Head Maralla Sialkot and Gujrat Districts, Punjab in quantitative terms, before the information is lost.

## Material and methods

### Study area description

Head Maralla is situated at the foot hills of Himalaya and lies almost between longitude 32°39’55.9 North and latitude 74°27’54.4 East at elevation of 233 masl. The Headwork was constructed in 1968 and is main water regulating structure at river Chenab. Map of different sites is shown in Figure 1, Map was selected and cartographic by GPS (Global Positioning System, Germany). There were four sites selected for collection of plants and observations; River Tavi (RT), Upstream Chenab (UC), River Manawarwala Tavi (RMT) and Bhalolpur side (BLP) (Figure 2). The climate of the area is warm and humid, further these sites are riverine, mostly wetland. Soil of the area is mostly sandy, while some area had loamy soil. During monsoon season, area is flooded by these three rivers Chenab, Tavi and Manawarwala.

**Figure 1.**
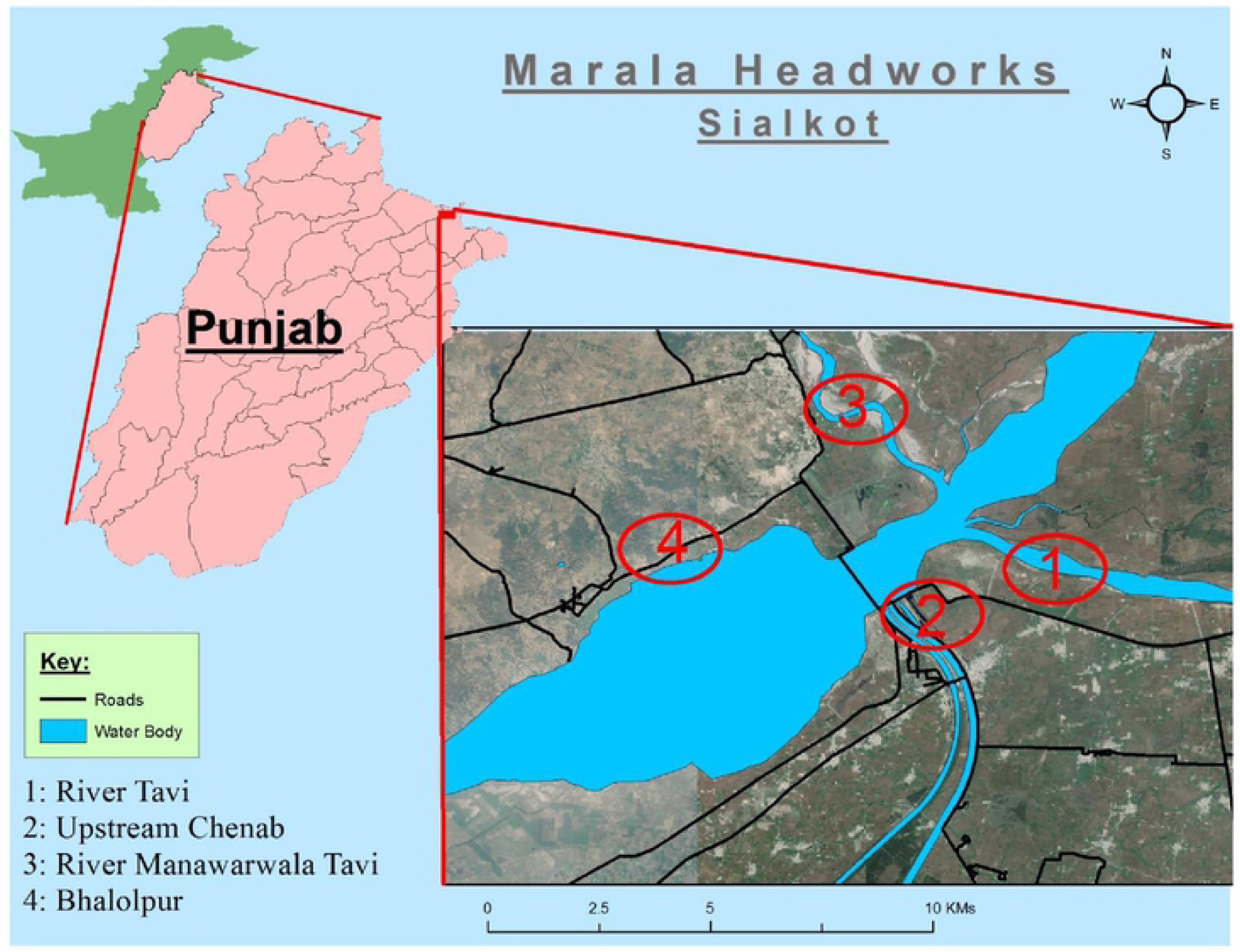
Map of Head Maralla showing four respective sites of the study area

**Figure 2.**
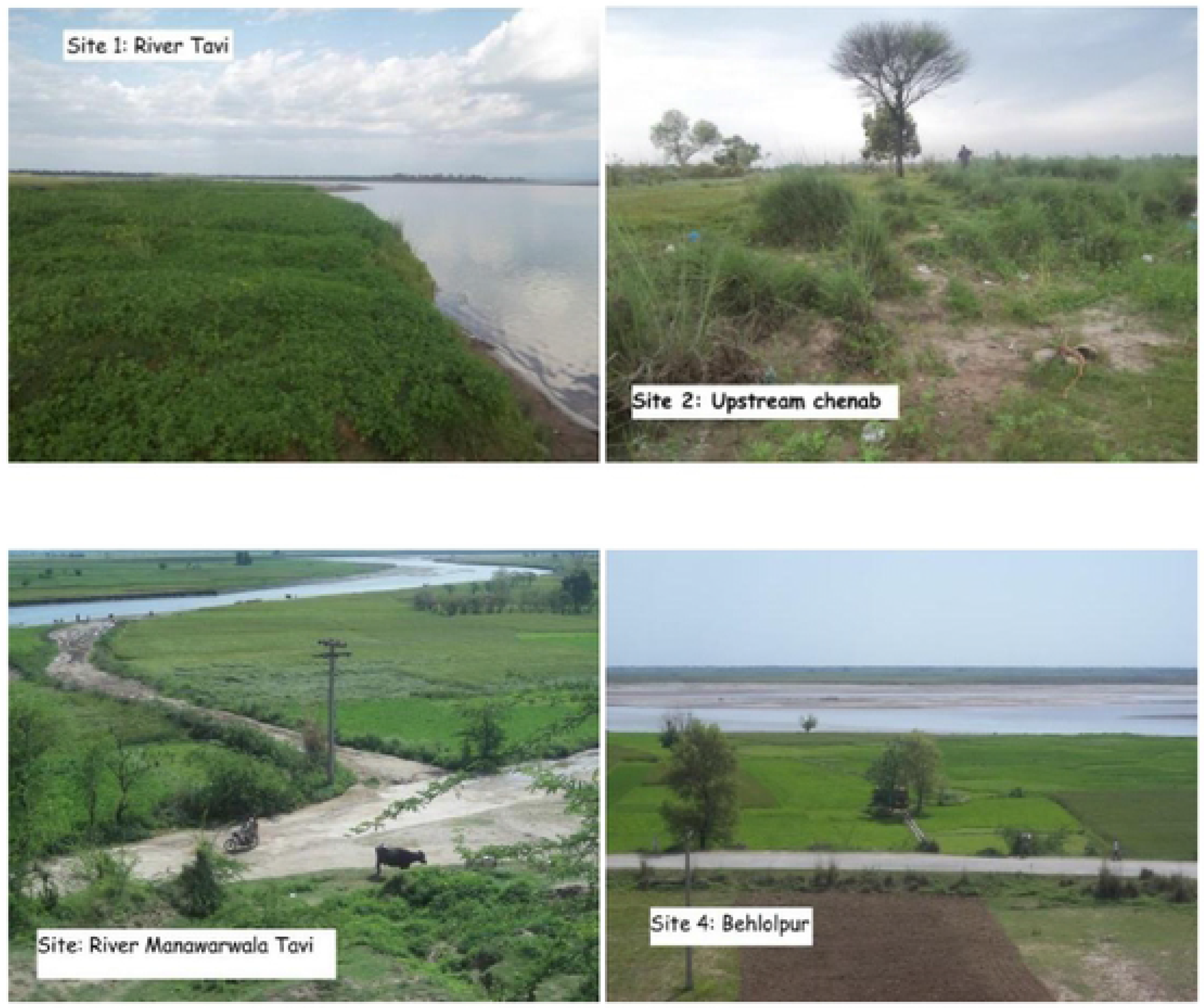
Four different sites of the study area are showing near the Head Maralla

### Demographic information

Study area is existed between two industrially big cities of the country i.e. Sialkot and Gujrat. City of Sialkot is situated in the north-east of the Punjab province. It is the 12th most populous city of the Pakistan and is well noted as entrepreneurial spirit and business climate, mentioned by ‘The Economist’. Goods including surgical instrumentation, sports, musical instrumentation and cutlery etc., constituted approximately $2 billion worth in 2018, nearly 10% of Pakistan’s total exports. Historically, Sialkot was of agricultural region with forests during Indus valley civilization while it had also been known as the winter capital of the State of Kashmir.

Sialkot District is surrounded by occupied Kashmir in the north east, Narowal District in the south east, Gujranwala and Sheikhpura Districts in the south west, Jummu district (India) in the north and Gujrat in the north west. It has four Tehsils Deska, Sambrial, Pasroor and Sialkot itself. The recorded history of Sialkot, a district of modern-day Pakistan, covers thousands of years. It has since its creation changed hands from Aryan, Persian, Greek, Hindu, Buddhist, Muslim, Sikh and British rule to the present-day federation of Pakistan. Sialkot district is spread over an area of 3,016 square kilometers [17, 18]. The district consisted of approximately 4.7 Million of population according to unpublished data of 6th population and housing census 2017 whereas 595,000 were mentioned by Demographia World Urban Areas Annual Edition [19, 20]. About 35% of the population constituted as urban.

Gujrat is an ancient city located between two main rivers Jhelum and Chenab. It is situated between Mirpur (North East), Jhelum (North West) and separates it from Jhelum district. On the east side, south east river Chenab is flowing, separating it from Gujranwala and Sialkot and in the west Mandi Bahauddin is present. It is administratively divided into four tehsils Kharian, Jalal Pur Jattan, Sarai Alamgir, and Gujrat itself [21]. District Gujrat is spread over an area of 3,192 square kilometers. Total population of Gujrat district is approximately 3.1 Million as unpublished data of 6th population and housing census 2017 reveals. Urban part of the population constituted as more than 37% females. Predominant language of the district is Punjabi, which according to the 1998 census is the first language of 98% of the population, while Urdu accounts for 1.1% [17, 18, 21].

Demographic information can be divided into following headings; a) climate and ecology of Sialkot, b) climate and ecology of Gujrat, c) ethnographic data, and d) data of agricultural land.

### Climate and ecology of District Sialkot

The geography and climate of the area is described as hot and humid during summer and cold in winter. May, June and July are the hottest months. During winter minimum temperature may drop to −2 °C and maximum ranged from 40 to 50 °C. The area is agriculture based with plain and fertile land. Monsoon season is the beauty of the area (mid of July to mid of August) where mid-September to mid-November remains hot during day time while nights cool down substantially. Low humidity prevails in these days. During winter from mid-November to March mild to warm weather can be observed and occasionally heavy rainfalls occurred. The average annual rainfall is 1000 mm [18, 19, 21].

### Climate and ecology of District Gujrat

This district has moderate climate, which is hot in summer and cold in winter. During peak summer, the day temperature shoots up to 50 °C, but the hot spells are comparatively shorter due to proximity of Azad Kashmir Mountains. The winter months are very pleasant and the minimum temperature may fall below 2 °C. The average rainfall on the Kashmir border is over 1000 mm, at Kharian it is 750 mm, at Gujrat 670 mm, and at Dinga 500 mm [21].

### Ethnographic data of Head Maralla

Head Maralla area is divided into two main religious communities, Muslim community in majority and Christians in minority. All ethnic groups use herbal medicines to cure various ailments. In Head Maralla, all of caste/tribes such as Jutt, Gujjar, Sheikh, Mughal, Nai, Faqir, Rajput, Malik, and Syed are present. According to local Government, estimated 5000 people lived around the Head Maralla, 70% people depends on agriculture. 15 hakims are engaged in different villages practicing herbal medicines. Local inhabitants who have traditional knowledge are about 40% (30% male) and (10% females). Average distribution of caste/tribe percentage is as; Jutt 40%, Gujjar 20%, Syed 10%, Mughal 10%, Malik 10%, Christian 5%, and Others are of 5% [17, 18].

#### Data of agricultural land

According to Agriculture Department, 450 hectares of land surrounding Head Maralla areas that is, the catchment area of Head Maralla along three rivers. 70% land is used for forest which is protected by army and under Forest Department while remaining land is used for agricultural purposes and highly influenced with anthropogenic activities. Irrigation Department facilitates irrigation system of Head Maralla. Two main canals in Head Maralla functioning; 1) Maralla Ravi link (water flow 22000 Cs), and 2) upper Chenab canal (water flow 16850 Cs). Old canal system constructed in 1905-1912 and construction of Head Maralla was accomplished during 1965-1968 while 66 Gates are present for regulation and balancing of water.

### Methodology for collecting data on plant diversity and ethnobotanical knowledge

Following methods were adapted to gather information; a) surveys, b) informant selection and ethnobotanical knowledge exploration, c) photography and inventory, d) preservation and taxonomical verification, e) botanical identification followed by f) quantitative analyses of ethnobotanical data.

#### Surveys of Head Maralla Areas

Six visits were arranged during summer and winter especially during monsoon season when maximum plant diversity were available and local inhabitants were consulted accordingly.

#### Informant selection and ethnobotanical knowledge exploration

Sites were selected after regular field visits and information was gathered about respondents and their expertise in traditional knowledge. Being local inhabitant, village men/women, army personnel and representative from irrigation and forest department were found to be well aware of these sites where there is a significant trend for utilization of traditional herbal recipes and other uses of wild plants in their daily routine for treating ailments and vice versa. In the respective area there is well awareness for cooperation and hospitality for students and faculty members, therefore after getting consent from irrigation and forest department there were no hurdle to meet people and get information for research purposes. In some villages local responsible person called as “Number Dar” who is the official bridge between government and villagers approached for effective community involvement and it is necessary to mention that without his assistance it was almost impossible to get knowledge in such kind of field surveys and studies. In case of women, their husbands and/or brothers were requested to assist, due to keeping societal norms because usually women are avoiding talking with strangers. Mean of communication was Punjabi and Urdu languages. Data were collected during spring, summer, monsoon, fall and winter depending on plant life cycles especially for flowering season. Ethnobotanical data and plants were collected from February 2014 to August 2017. Local respondents were contacted during spring and summer when the floristic diversity was on peak that means before monsoon season which bring high amount of water, even flood in all of the adjacent areas.

Along local inhabitants, many hakims, practioners and healers were also consulted and discussed about various plants, their collection methods, recipes, formulation, trade, their availability and multiplication and even preservation. Rapid appraisal approach (RAA) was conducted to collect the indigenous knowledge. Survey was based on direct interaction with indigenous people through group discussions, corner meetings and semi-structured interviews following the method by Martin [22]. Hundred local informants were contacted formally to obtain information, in which informants were belonged to different age groups i-e., 40-50, 51-60, 61-70, 71-80 and >80 years old. Out of 100, 73 were male and 27 were female. While 15 were local healers (Hakims), others were affiliated with agriculture, fish hunting, house wives, local government, forest, irrigation and agriculture departments’ employee by professional (Table 1). Although different age group people had knowledge but elders were proved more authentic and had command in traditional knowledge, interestingly the other local people know such prominent elders and advised to contact them. To ensure the strong validity of traditional knowledge, a continuous relationship was maintained with the local people throughout survey.

**Table 1.** Demographic data of informants recorded at Head Maralla

| Age group | Informants | Male | Percentage | Female | Percentage |
| --- | --- | --- | --- | --- | --- |
| 40-50 | 14 | 10 | 13.69 | 04 | 14.81 |

| 51-60 | 22 | 15 | 20.54 | 07 | 29.97 |
| --- | --- | --- | --- | --- | --- |
| 61-70 | 33 | 24 | 32.87 | 09 | 33.31 |
| 71-80 | 26 | 20 | 27.39 | 06 | 27.22 |
| >80 | 05 | 04 | 5.47 | 01 | 3.70 |
| **Total** | **100** | **73** |  | **27** |  |
| **Educational background** |  | **levels** | **Informants** |  |  |
| Illiterate or primary pass (5 years) |  | School | 42 |  |  |
| Middle pass (8 years) |  | Middle School | 07 |  |  |
| Matric (10 years) |  | High School | 21 |  |  |
| Undergraduate (14 years) |  | College | 21 |  |  |
| Graduate (16 years) or above |  | University | 09 |  |  |

Interviews were conducted after obtaining informed consent (IC) from the interviewees. There were two parts of the questionnaire a) demographic data and b) uses of plants. Various applications were then divided into medicinal and industrial plants. The traditional knowledge was documented as inventory which was consisted of local name of the plants, parts used, methods of preparation, mode of usage, and the diseases treated (questionnaire as supplementary file).

#### Photography and Inventory

Photography of different sites, vegetation and some unique plants of the sites were captured by cameras ‘Cannon d3000’ and ‘Sony cyber shot’ and also to get assistance for identification. An inventory was established after comparing the data with available literature. Information associated with local culture, local norms and flora, avian fauna and general information for population was also documented.

#### Preservation and taxonomical verifications

Collected specimens were assigned voucher numbers and stored in the newly established Herbarium of Department of Botany, University of Gujrat, Gujrat, Pakistan as per following the known methods of drying, pressing and preserving i.e., the plants were pressed for dryness, poisoned (1% HgCl_2_ sol.) and then mounted on herbarium sheets. Collected species were identified and compared with Flora of Pakistan along accessible literature while complicated specimens were verified with the collection available at Quaid-i-Azam University, Islamabad, Pakistan [23].

#### Botanical Identification

During field surveys, identification was mainly based on the local names of plants, with the help of local informants. For taxonomic confirmation, the Flora of Pakistan (http://www.efloras.org/index.aspx) was followed. Further proper classification and names were cross checked with APG IV (where applicable), USDA plant database and ‘The Plant List’. In the current studies majority of the names are according to ‘The Plant List’ for correct botanical name confirmation (http://www.theplantlist.org).

### Quantitative Analyses of Ethnobotanical data

Quantitative indices, including Informant Consensus Factor (ICF) and Use Value (UV) were applied, as following;

#### Informant’s Consensus Factor

To evaluate the use of medicinal plants and their finding for bioactive compounds the most widely used tool, Informants consensus factor (*Fic*) was calculated [24]. This technique was used to find/give more prominence of the information regarding particular types of ailment categories. This tool estimates relationship between the “number of used reports in each category *(Nur)* minus the number of specie used *(nt)”*and the “number of used-reports in each category minus 1”. While the low value close to 0 represents the plants have been chosen randomly for a few or a single disease or that informant did not shared knowledge about the use of plants [25]. FIC can be calculated using the following formula

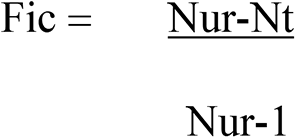

#### Use values

During interview information of different uses were obtained, the use value (UV) of the plant species has been calculated, using the following formula [26];

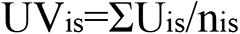

Where UV_is_ = the use value of the species ‘s’ mentioned by the informant ‘i’; ΣU_is_ = the number of uses of species ‘s’ mentioned in each event by the informant ‘I’; n_is_= the number of events in which the informant represented by ‘i’ and cited species by s. UVs were higher if there were many use reports of a plant, implying that the plant is important, whereas they were near zero if there were few reports related to its use.

#### Comparison with neighboring areas

A comprehensive analysis has been conducted for comparing current findings with previous published data. 40 different studies were found most suitable to provide comparison about area, study year, families, total species common in both areas, plants with similar uses (%), and plants with dissimilar uses (%), species enlisted in specified area, Jaccard index and citation.

## 1. Results

### Plant diversity of Head Maralla

One hundred nineteen (119) plant species belonging to 54 families were recorded at Head Maralla are shown to assess their occurrence and diversity in Figure 3. It reflects that 88 species of dicot, 20 monocot, 05 pteridophytes, 04 bryophytes and 02 algae are hereby documented. Lakes, ponds and ditches present in the area were also studied in order to measure aquatic flora and their medicinal uses. *Nelumbium, Typha* and *Ceratophyllum* were most important genera found from the water bodies. At some places, it has been noted that *Typha* is abundantly used by the local peoples for various purposes. Among bryophytes *Marchantia polymorpha* L.*, Riccia cavernosa* Hoffm*., Polytricum* sp., and *Funaria hygrometrica* Hedw., were common species while in case of ferns, *Osmunda regalis* L., was mostly common. Inventory of floral diversity was established and provided (Table 2). Plants were collected in different seasons as pre-monsoon, monsoon and post-monsoon. Moreover, they were observed at both terrestrial and aquatic habitat. Presence of common genera including *Ipomoea, Euphorbia, Ficus, Solanum, Acacia* and *Amaranthus* were recorded at all sites.

**Figure 3.**
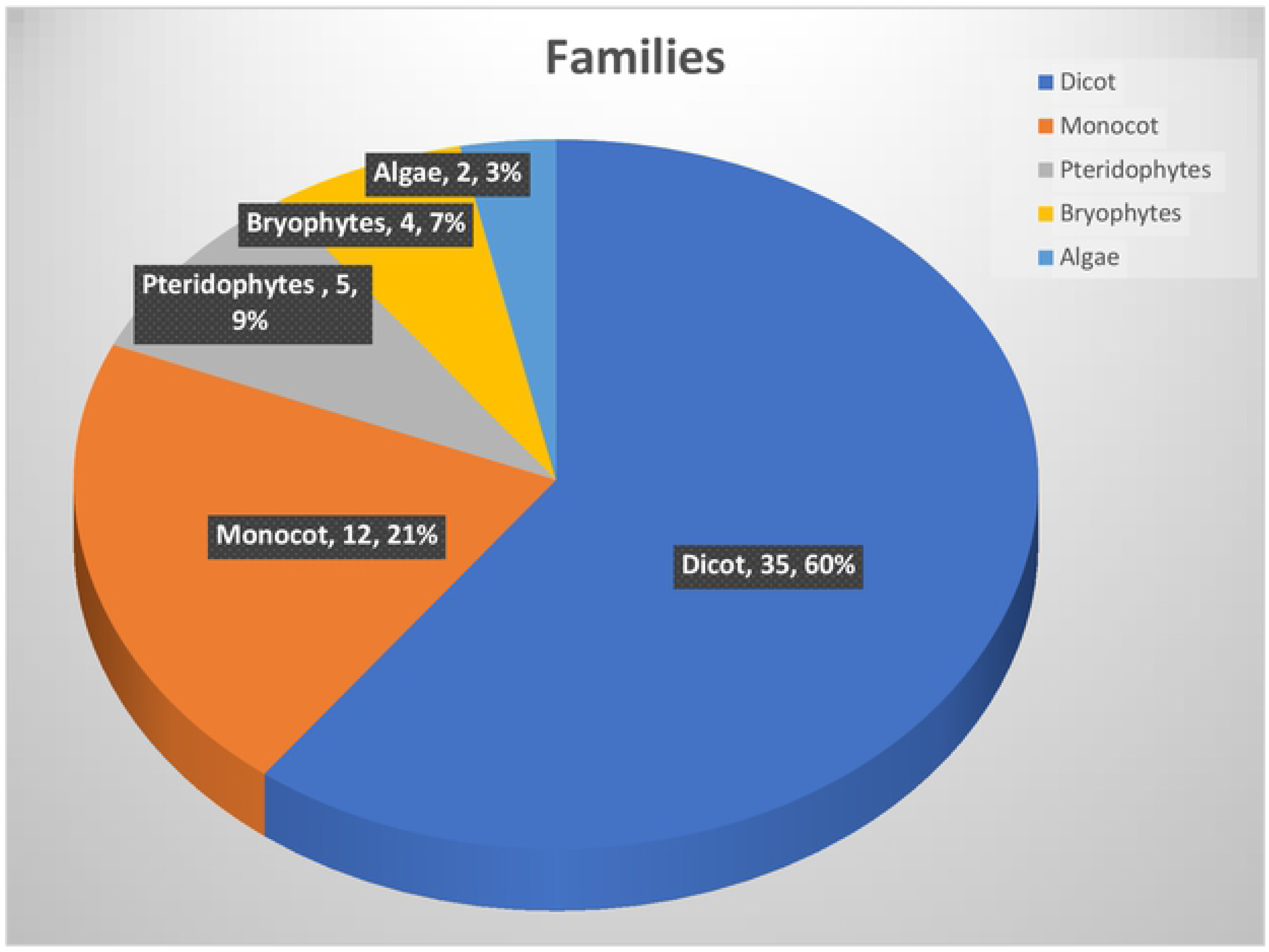
Summary of floral diversity recorded at Head Maralla

**Table 2.**
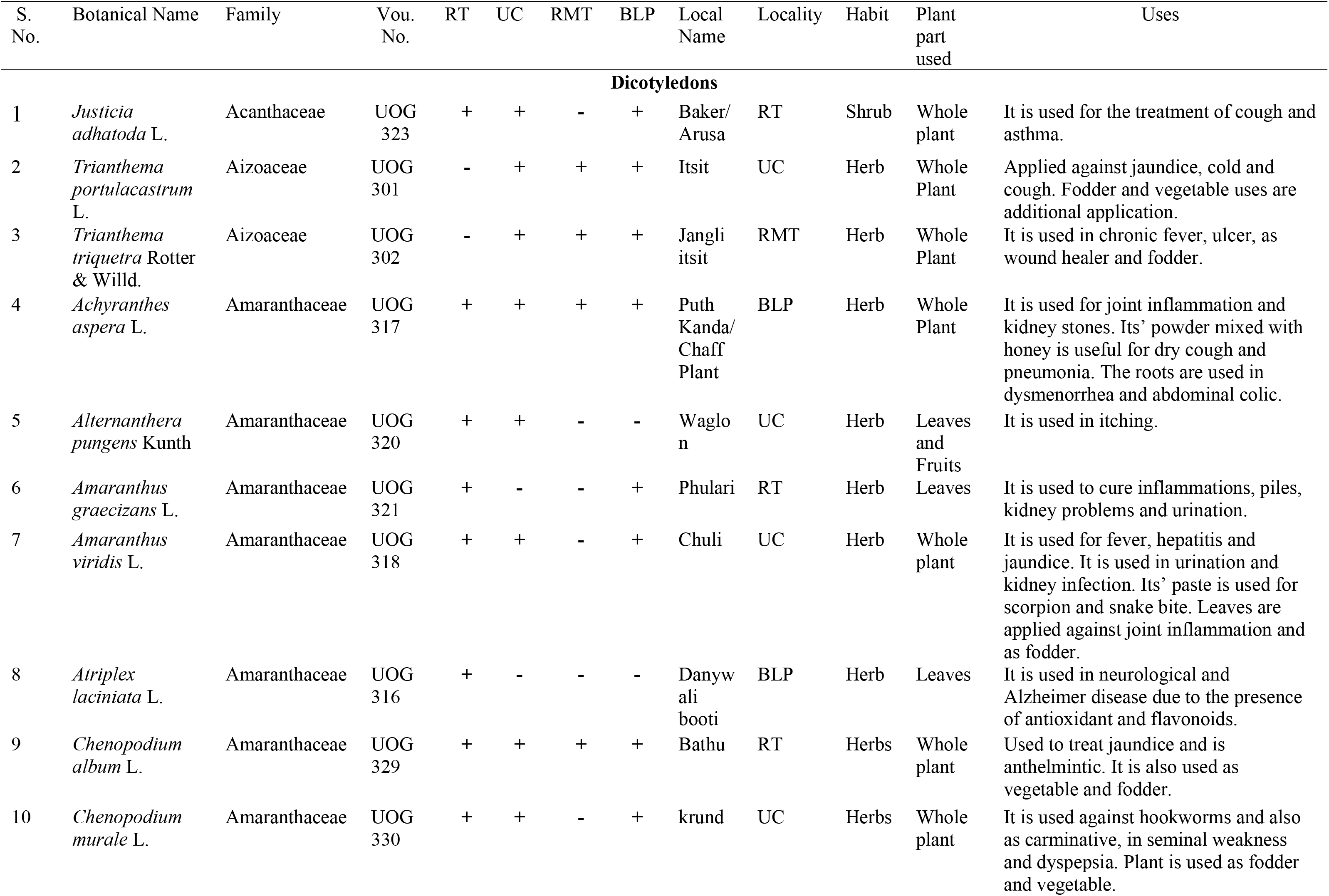
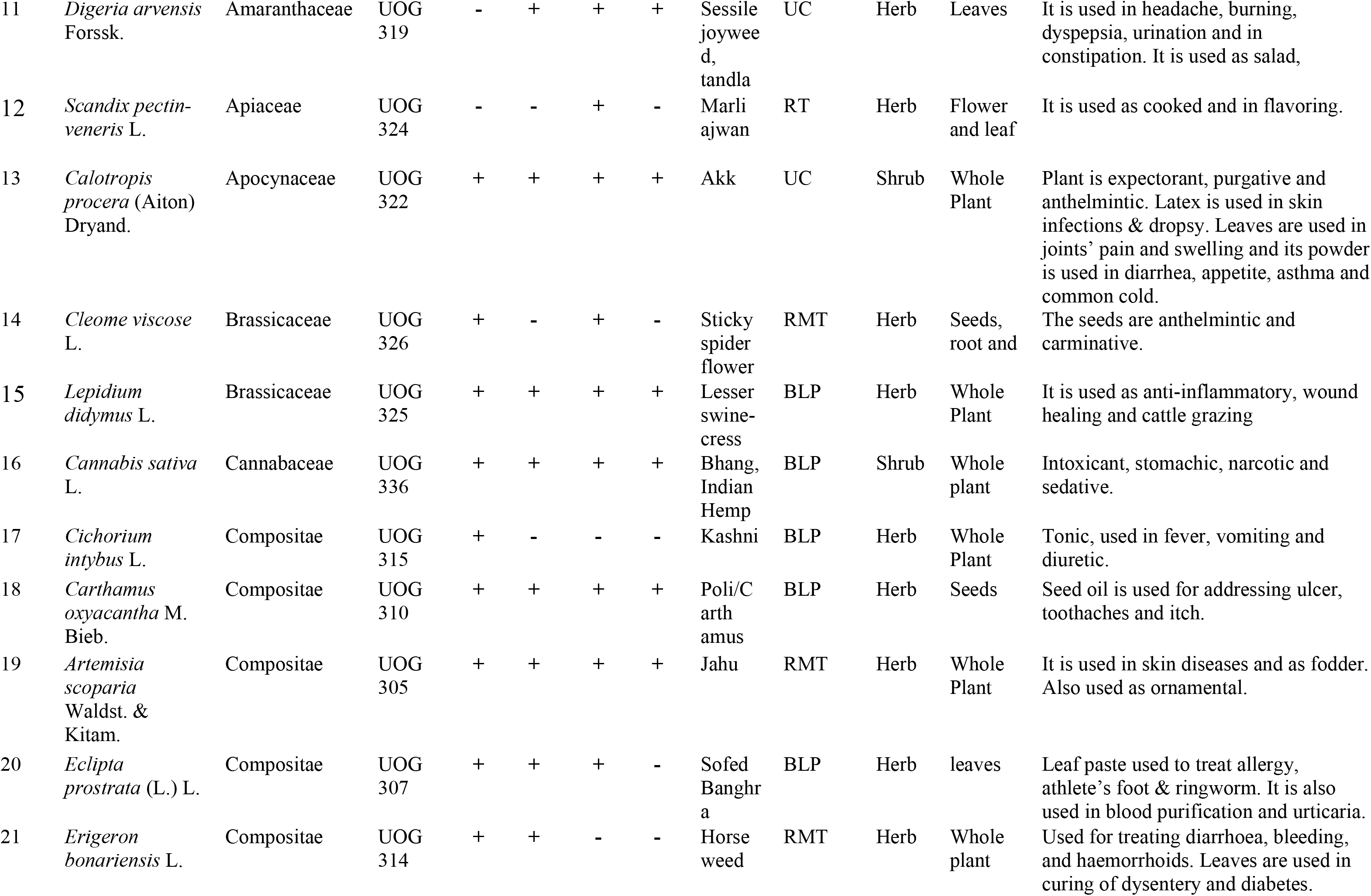
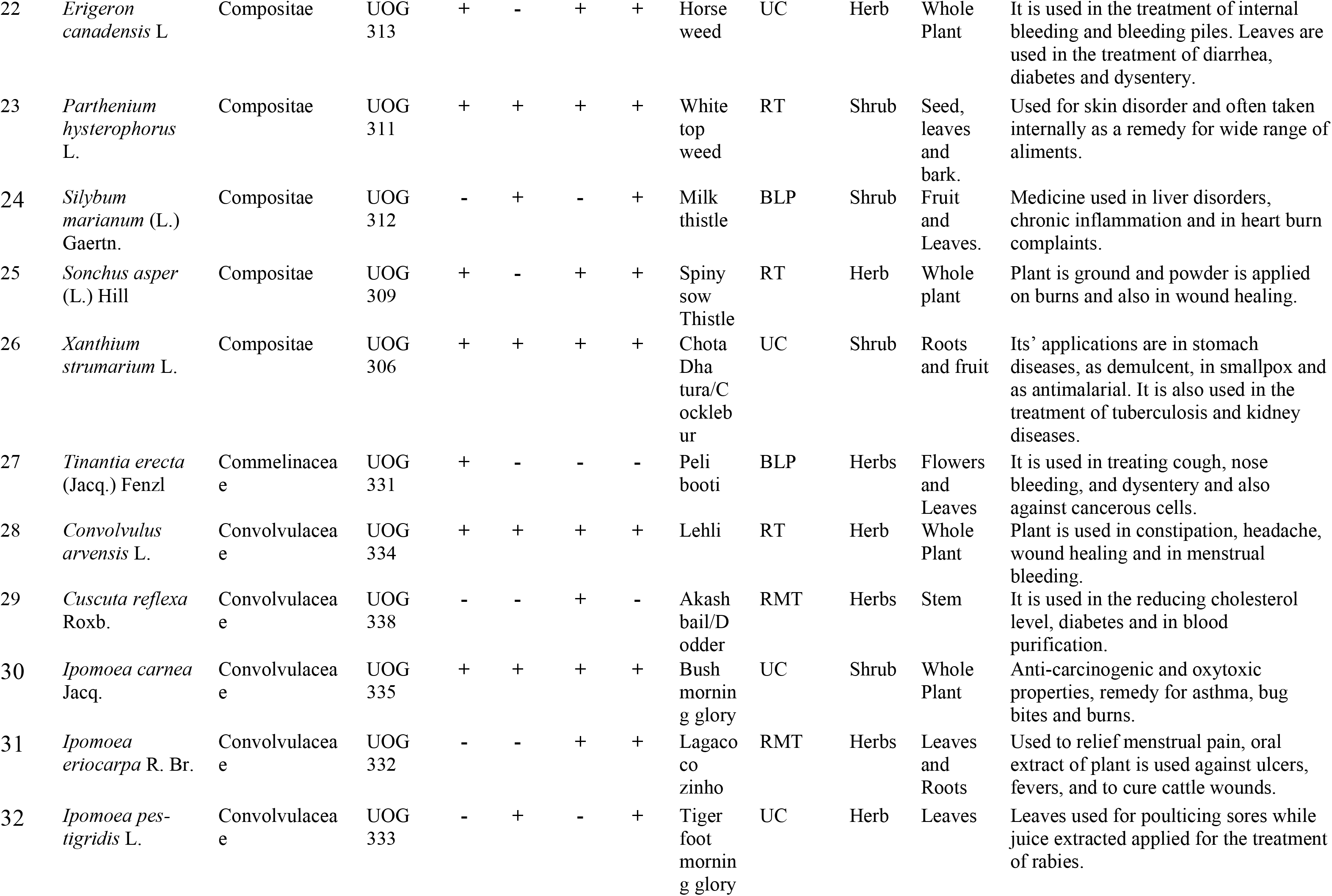
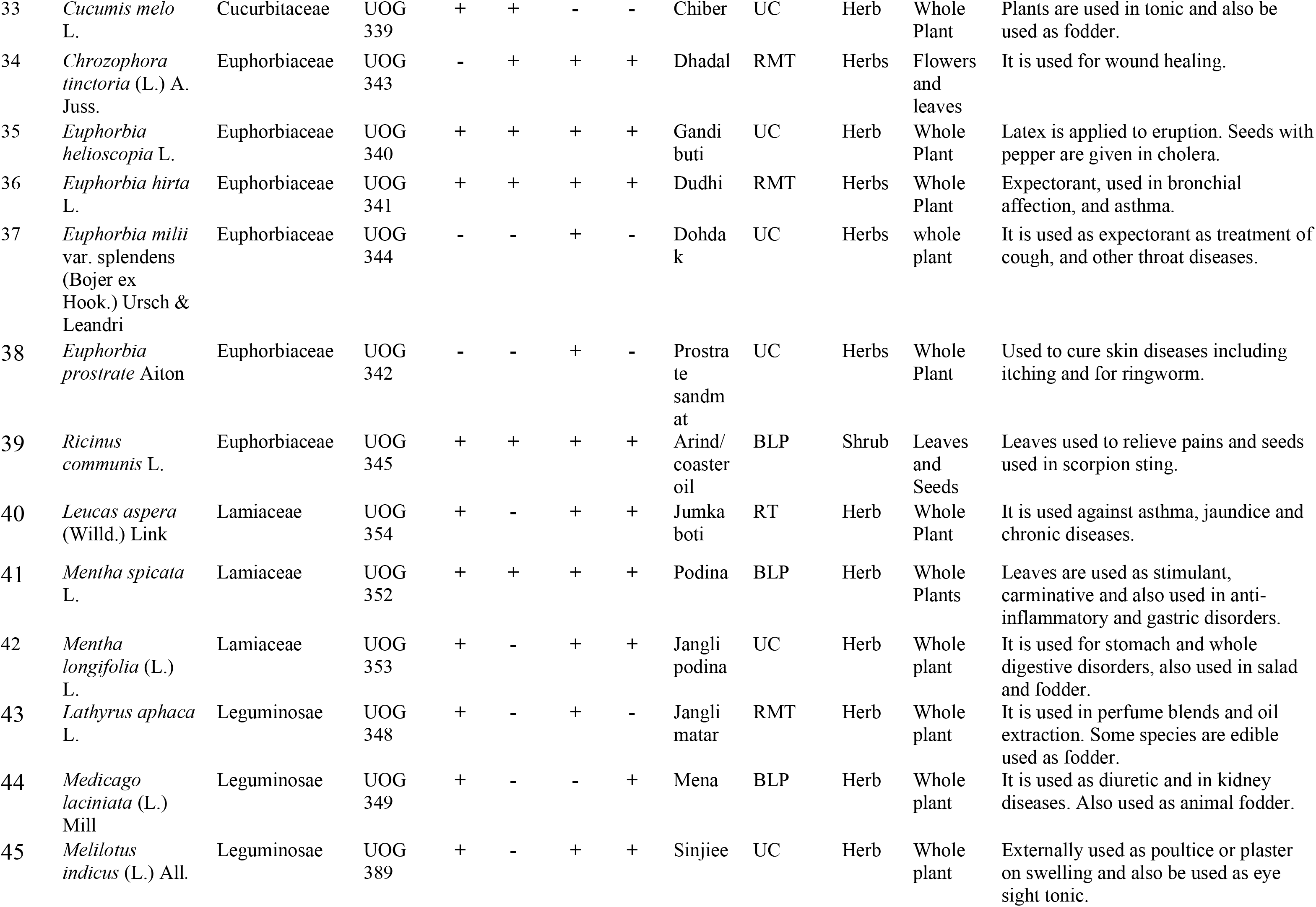
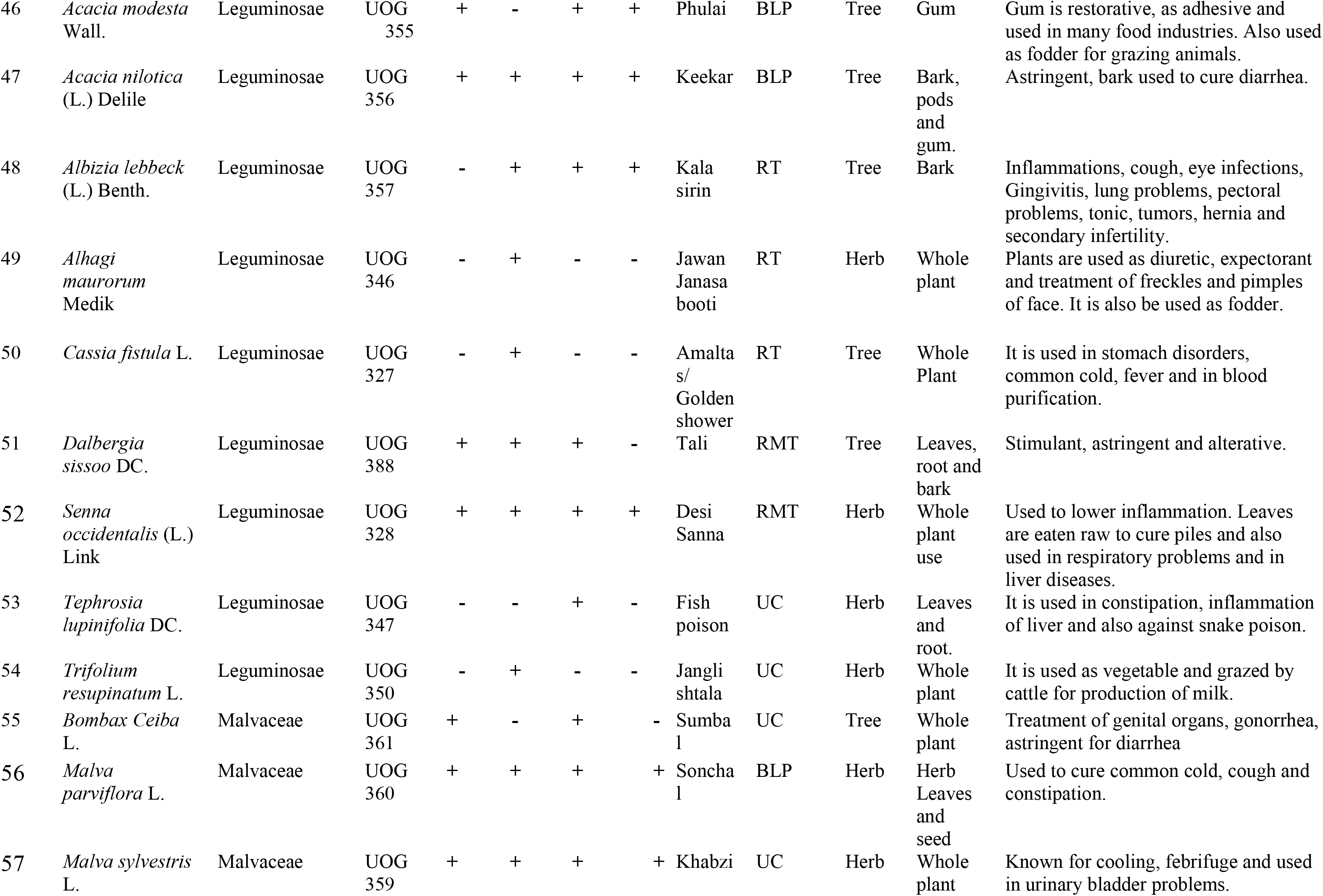
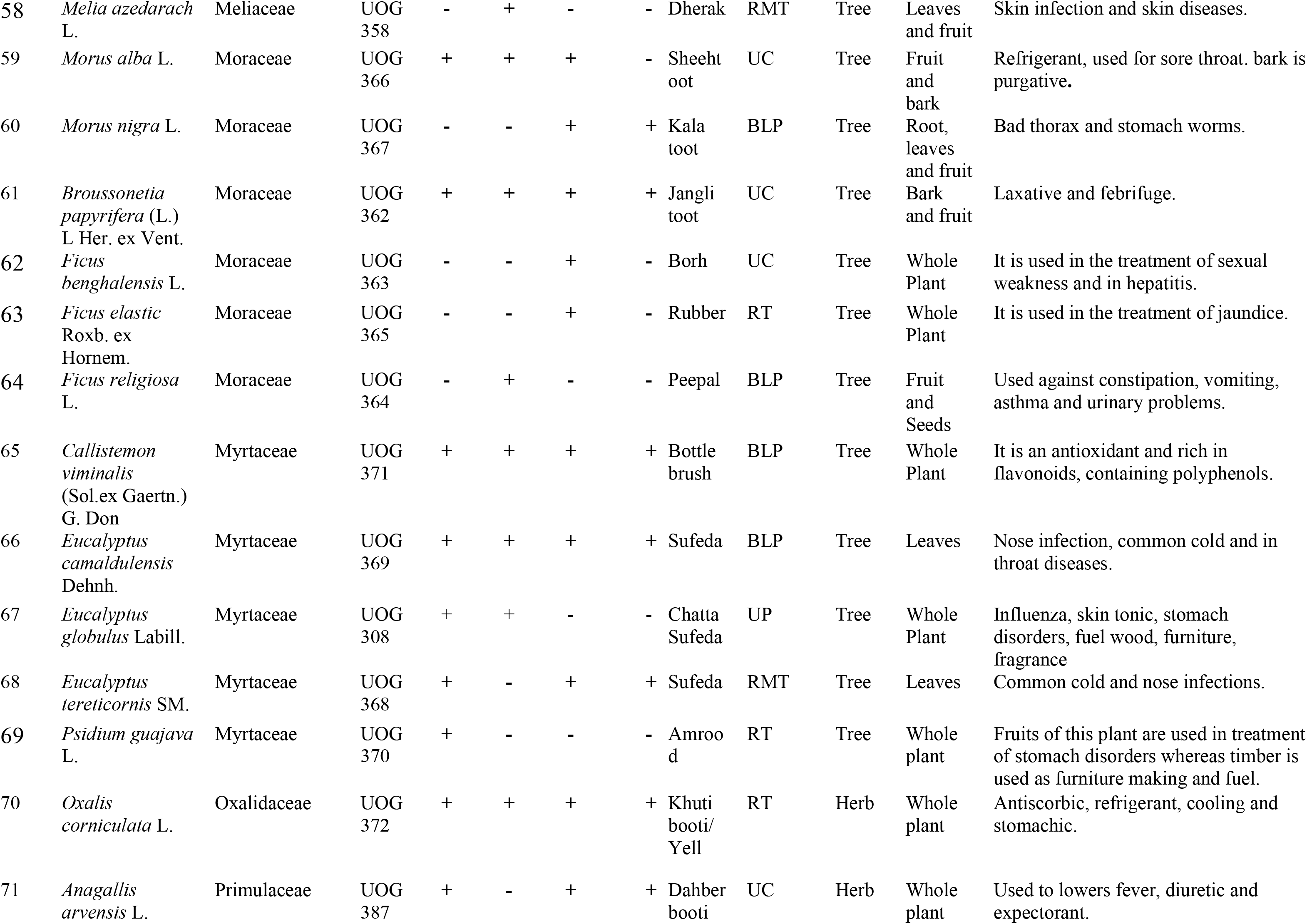
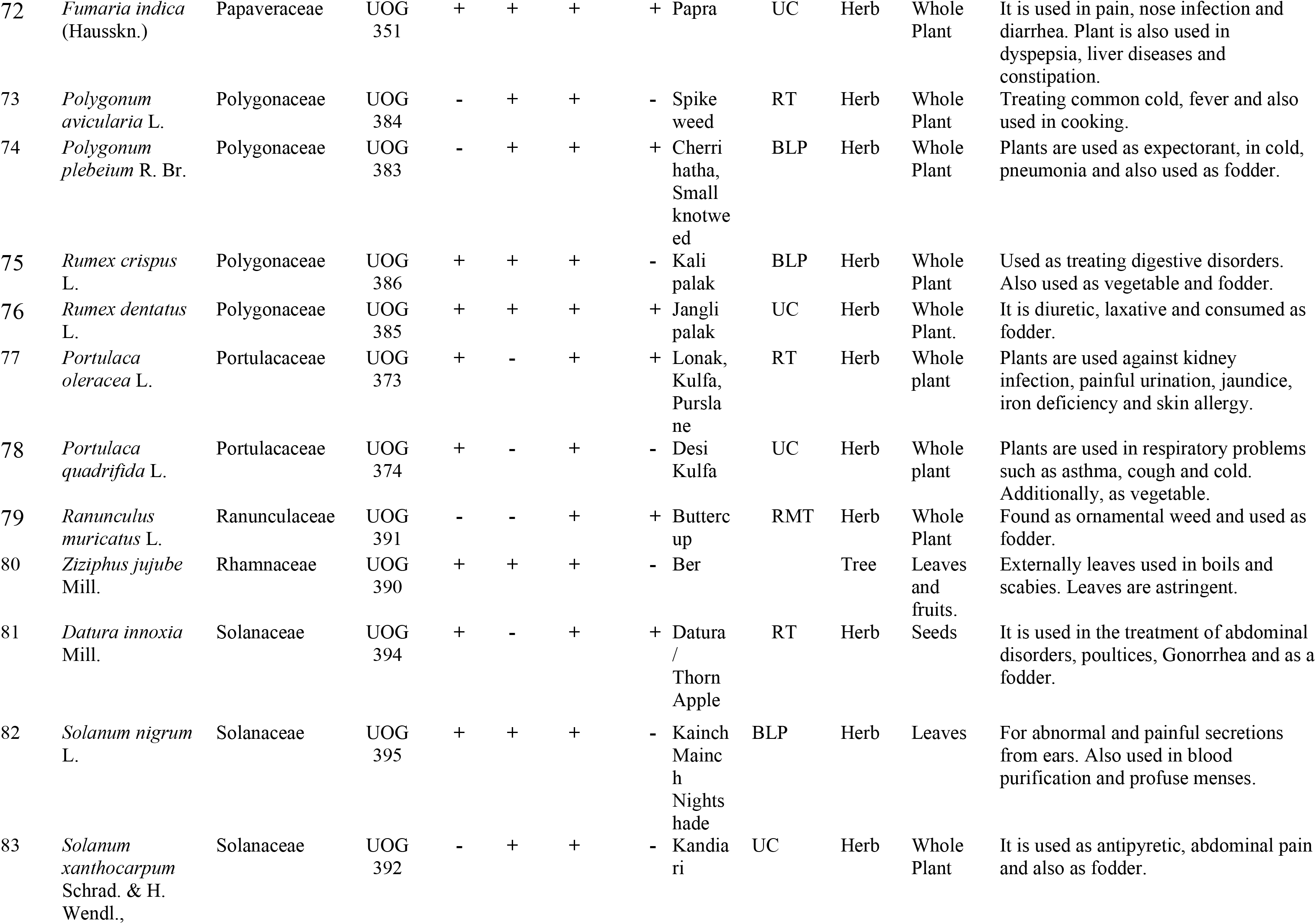
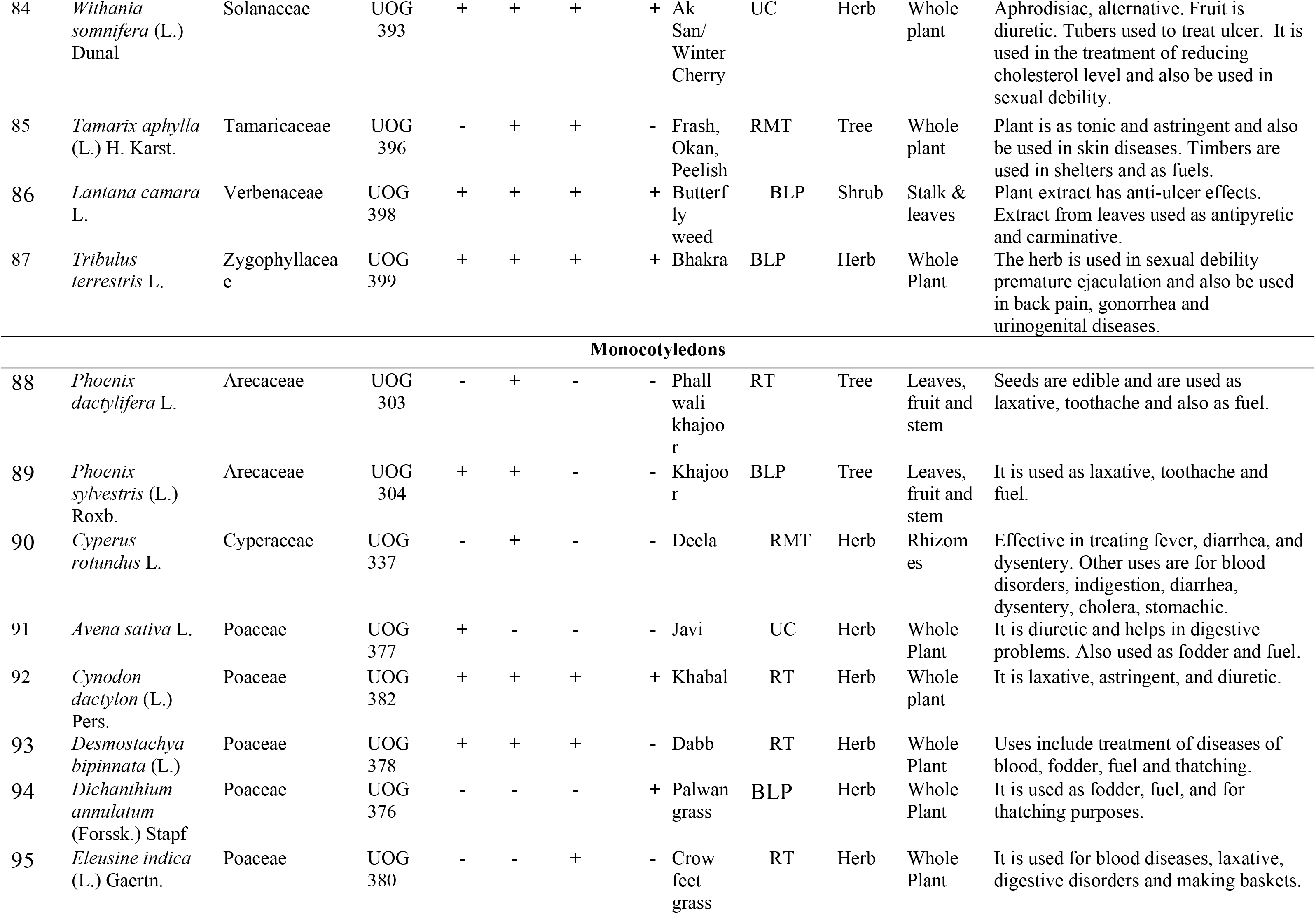
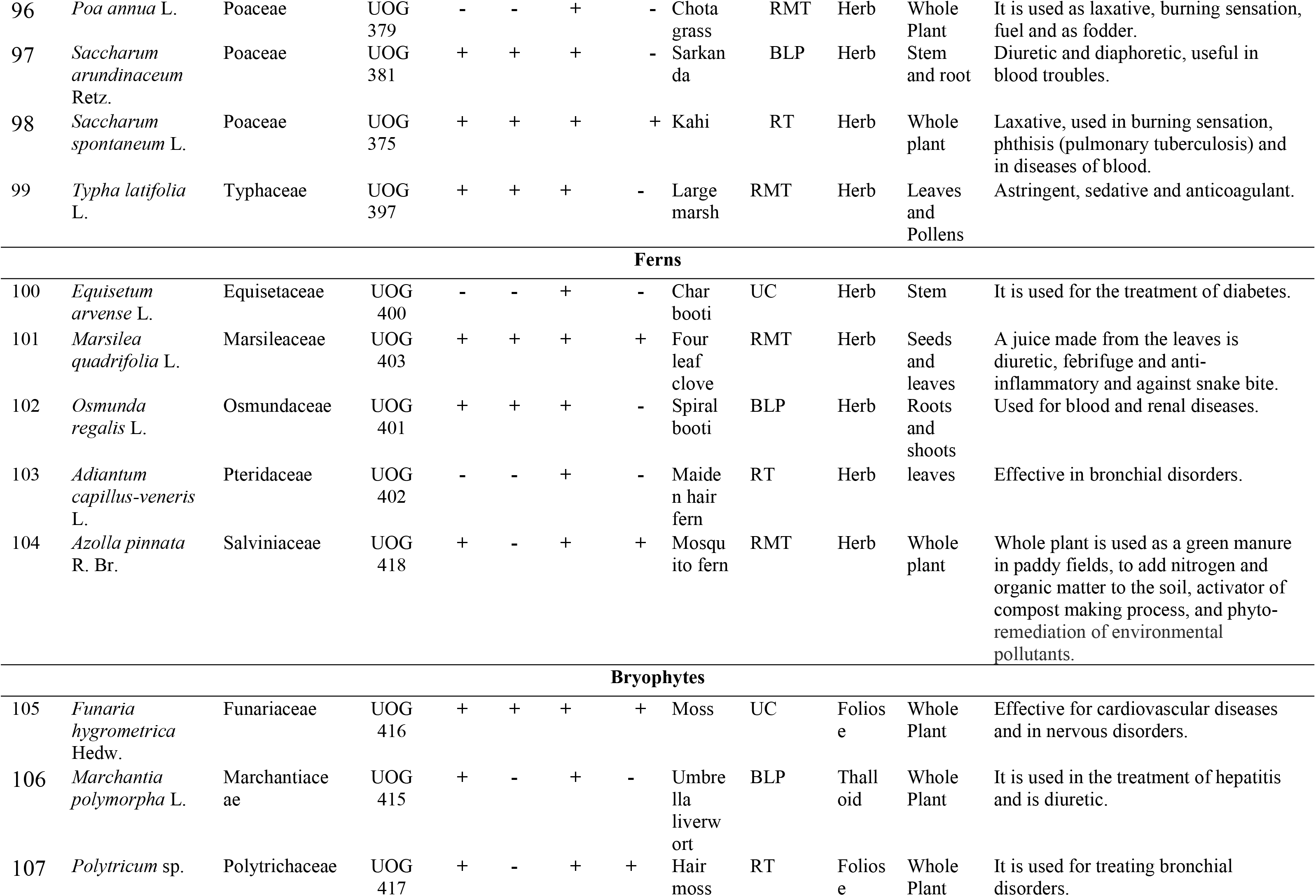
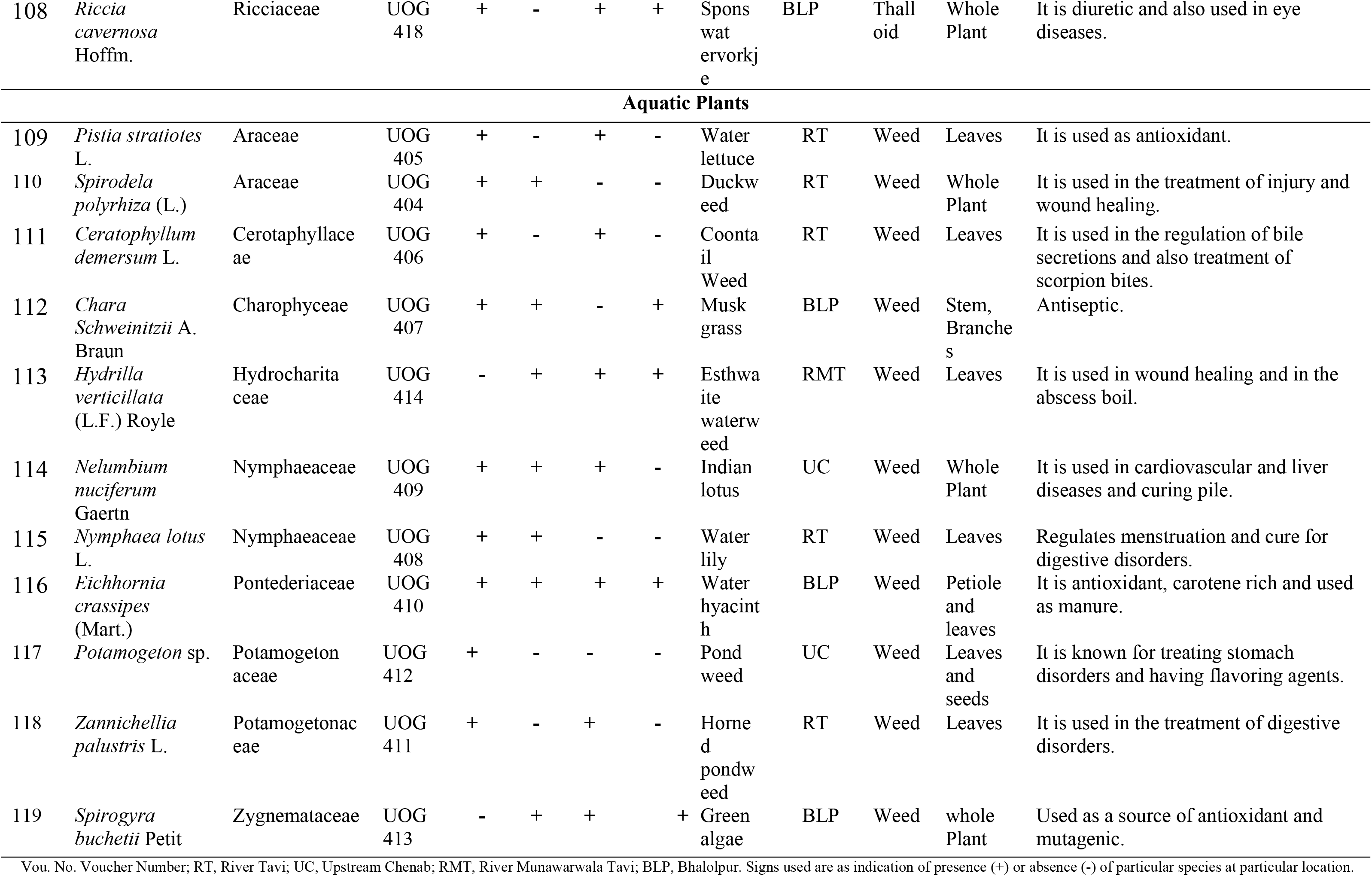
Inventory of wild plants with medicinal and industrial uses along with locality reported from Head Maralla

| S. No. | Botanical Name | Family | Vou. No. | RT | UC | RMT | BLP | Local Name | Locality | Habit | Plant part used | Uses |
| --- | --- | --- | --- | --- | --- | --- | --- | --- | --- | --- | --- | --- |
| **Dicotyledons** |  |  |  |  |  |  |  |  |  |  |  |  |
| 1 | *Justicia adhatoda* L. | Acanthaceae | UOG 323 | + | + | - | + | Baker/ Arusa | RT | Shrub | Whole plant | It is used for the treatment of cough and asthma. |
| 2 | *Trianthema portulacastrum* L. | Aizoaceae | UOG 301 | - | + | + | + | Itsit | UC | Herb | Whole Plant | Applied against jaundice, cold and cough. Fodder and vegetable uses are additional application. |
| 3 | *Trianthema triquetra* Rotter & Willd. | Aizoaceae | UOG 302 | - | + | + | + | Jangli itsit | RMT | Herb | Whole Plant | It is used in chronic fever, ulcer, as wound healer and fodder. |
| 4 | *Achyranthes aspera* L. | Amaranthaceae | UOG 317 | + | + | + | + | Puth Kanda/ Chaff Plant | BLP | Herb | Whole Plant | It is used for joint inflammation and kidney stones. Its' powder mixed with honey is useful for dry cough and pneumonia. The roots are used in dysmenorrhea and abdominal colic. |
| 5 | *Alternanthera pungens* Kunth | Amaranthaceae | UOG 320 | + | + | - | - | Waglon | UC | Herb | Leaves and Fruits | It is used in itching. |
| 6 | *Amaranthus graecizans* L. | Amaranthaceae | UOG 321 | + | - | - | + | Phulari | RT | Herb | Leaves | It is used to cure inflammations, piles, kidney problems and urination. |
| 7 | *Amaranthus viridis* L. | Amaranthaceae | UOG 318 | + | + | - | + | Chuli | UC | Herb | Whole plant | It is used for fever, hepatitis and jaundice. It is used in urination and kidney infection. Its' paste is used for scorpion and snake bite. Leaves are applied against joint inflammation and as fodder. |
| 8 | *Atriplex laciniata* L. | Amaranthaceae | UOG 316 | + | - | - | - | Danywali booti | BLP | Herb | Leaves | It is used in neurological and Alzheimer disease due to the presence of antioxidant and flavonoids. |
| 9 | *Chenopodium album* L. | Amaranthaceae | UOG 329 | + | + | + | + | Bathu | RT | Herbs | Whole plant | Used to treat jaundice and is anthelmintic. It is also used as vegetable and fodder. |
| 10 | *Chenopodium murale* L. | Amaranthaceae | UOG 330 | + | + | - | + | krund | UC | Herbs | Whole plant | It is used against hookworms and also as carminative, in seminal weakness and dyspepsia. Plant is used as fodder and vegetable. |
| 11 | *Digeria arvensis* Forssk. | Amaranthaceae | UOG 319 | - | + | + | + | Sessile joyweed, tandla | UC | Herb | Leaves | It is used in headache, burning, dyspepsia, urination and in constipation. It is used as salad, |
| 12 | *Scandix pecten-veneris* L. | Apiaceae | UOG 324 | - | - | + | - | Marli ajwan | RT | Herb | Flower and leaf | It is used as cooked and in flavoring. |
| 13 | *Calotropis procera* (Aiton) Dryand. | Apocynaceae | UOG 322 | + | + | + | + | Akk | UC | Shrub | Whole Plant | Plant is expectorant, purgative and anthelmintic. Latex is used in skin infections & dropsy. Leaves are used in joints' pain and swelling and its powder is used in diarrhea, appetite, asthma and common cold. |
| 14 | *Cleome viscosa* L. | Brassicaceae | UOG 326 | + | - | + | - | Sticky spider flower | RMT | Herb | Seeds, root and | The seeds are anthelmintic and carminative. |
| 15 | *Lepidium didymus* L. | Brassicaceae | UOG 325 | + | + | + | + | Lesser swine-cress | BLP | Herb | Whole Plant | It is used as anti-inflammatory, wound healing and cattle grazing |
| 16 | *Cannabis sativa* L. | Cannabaceae | UOG 336 | + | + | + | + | Bhang, Indian Hemp | BLP | Shrub | Whole plant | Intoxicant, stomachic, narcotic and sedative. |
| 17 | *Cichorium intybus* L. | Compositae | UOG 315 | + | - | - | - | Kashni | BLP | Herb | Whole Plant | Tonic, used in fever, vomiting and diuretic. |
| 18 | *Carthamus oxyacantha* M. Bieb. | Compositae | UOG 310 | + | + | + | + | Poli/C arthamus | BLP | Herb | Seeds | Seed oil is used for addressing ulcer, toothaches and itch. |
| 19 | *Artemisia scoparia* Waldst. & Kitam. | Compositae | UOG 305 | + | + | + | + | Jahu | RMT | Herb | Whole Plant | It is used in skin diseases and as fodder. Also used as ornamental. |
| 20 | *Eclipta prostrata* (L.) L. | Compositae | UOG 307 | + | + | + | - | Sofed Banghra | BLP | Herb | leaves | Leaf paste used to treat allergy, athlete's foot & ringworm. It is also used in blood purification and urticaria. |
| 21 | *Erigeron bonariensis* L. | Compositae | UOG 314 | + | + | - | - | Horse weed | RMT | Herb | Whole plant | Used for treating diarrhoea, bleeding, and haemorrhoids. Leaves are used in curing of dysentery and diabetes. |
| 22 | *Erigeron canadensis* L. | Compositae | UOG 313 | + | - | + | + | Horse weed | UC | Herb | Whole Plant | It is used in the treatment of internal bleeding and bleeding piles. Leaves are used in the treatment of diarrhea, diabetes and dysentery. |
| 23 | *Parthenium hysterophorus* L. | Compositae | UOG 311 | + | + | + | + | White top weed | RT | Shrub | Seed, leaves and bark. | Used for skin disorder and often taken internally as a remedy for wide range of ailments. |
| 24 | *Silybum marianum* (L.) Gaertn. | Compositae | UOG 312 | - | + | - | + | Milk thistle | BLP | Shrub | Fruit and Leaves. | Medicine used in liver disorders, chronic inflammation and in heart burn complaints. |
| 25 | *Sonchus asper* (L.) Hill | Compositae | UOG 309 | + | - | + | + | Spiny sow Thistle | RT | Herb | Whole plant | Plant is ground and powder is applied on burns and also in wound healing. |
| 26 | *Xanthium strumarium* L. | Compositae | UOG 306 | + | + | + | + | Chota Dha tura/Cocklebur | UC | Shrub | Roots and fruit | Its' applications are in stomach diseases, as demulcent, in smallpox and as antimalarial. It is also used in the treatment of tuberculosis and kidney diseases. |
| 27 | *Tinantia erecta* (Jacq.) Fenzl | Commelinaceae | UOG 331 | + | - | - | - | Peli booti | BLP | Herbs | Flowers and Leaves | It is used in treating cough, nose bleeding, and dysentery and also against cancerous cells. |
| 28 | *Convolvulus arvensis* L. | Convolvulaceae | UOG 334 | + | + | + | + | Lehli | RT | Herb | Whole Plant | Plant is used in constipation, headache, wound healing and in menstrual bleeding. |
| 29 | *Cuscuta reflexa* Roxb. | Convolvulaceae | UOG 338 | - | - | + | - | Akash bail/Dodder | RMT | Herbs | Stem | It is used in the reducing cholesterol level, diabetes and in blood purification. |
| 30 | *Ipomoea carnea* Jacq. | Convolvulaceae | UOG 335 | + | + | + | + | Bush morning glory | UC | Shrub | Whole Plant | Anti-carcinogenic and oxytoxic properties, remedy for asthma, bug bites and burns. |
| 31 | *Ipomoea eriocarpa* R. Br. | Convolvulaceae | UOG 332 | - | - | + | + | Lagaco zinho | RMT | Herbs | Leaves and Roots | Used to relief menstrual pain, oral extract of plant is used against ulcers, fevers, and to cure cattle wounds. |
| 32 | *Ipomoea pes-tigridis* L. | Convolvulaceae | UOG 333 | - | + | - | + | Tiger foot morning glory | UC | Herb | Leaves | Leaves used for poulticing sores while juice extracted applied for the treatment of rabies. |
| 33 | *Cucumis melo* L. | Cucurbitaceae | UOG 339 | + | + | - | - | Chiber | UC | Herb | Whole Plant | Plants are used in tonic and also be used as fodder. |
| 34 | *Chrozophora tinctoria* (L.) A. Juss. | Euphorbiaceae | UOG 343 | - | + | + | + | Dhadal | RMT | Herbs | Flowers and leaves | It is used for wound healing. |
| 35 | *Euphorbia helioscopia* L. | Euphorbiaceae | UOG 340 | + | + | + | + | Gandi buti | UC | Herb | Whole Plant | Latex is applied to eruption. Seeds with pepper are given in cholera. |
| 36 | *Euphorbia hirta* L. | Euphorbiaceae | UOG 341 | + | + | + | + | Dudhi | RMT | Herbs | Whole Plant | Expectorant, used in bronchial affection, and asthma. |
| 37 | *Euphorbia milii* var. splendens (Bojer ex Hook.) Ursch & Leandri | Euphorbiaceae | UOG 344 | - | - | + | - | Dohda k | UC | Herbs | whole plant | It is used as expectorant as treatment of cough, and other throat diseases. |
| 38 | *Euphorbia prostrata* Aiton | Euphorbiaceae | UOG 342 | - | - | + | - | Prostrate sandm at | UC | Herbs | Whole Plant | Used to cure skin diseases including itching and for ringworm. |
| 39 | *Ricinus communis* L. | Euphorbiaceae | UOG 345 | + | + | + | + | Arind/coaster oil | BLP | Shrub | Leaves and Seeds | Leaves used to relieve pains and seeds used in scorpion sting. |
| 40 | *Leucas aspera* (Willd.) Link | Lamiaceae | UOG 354 | + | - | + | + | Jumka boti | RT | Herb | Whole Plant | It is used against asthma, jaundice and chronic diseases. |
| 41 | *Mentha spicata* L. | Lamiaceae | UOG 352 | + | + | + | + | Podina | BLP | Herb | Whole Plants | Leaves are used as stimulant, carminative and also used in anti-inflammatory and gastric disorders. |
| 42 | *Mentha longifolia* (L.) L. | Lamiaceae | UOG 353 | + | - | + | + | Jangli podina | UC | Herb | Whole plant | It is used for stomach and whole digestive disorders, also used in salad and fodder. |
| 43 | *Lathyrus aphaca* L. | Leguminosae | UOG 348 | + | - | + | - | Jangli matar | RMT | Herb | Whole plant | It is used in perfume blends and oil extraction. Some species are edible used as fodder. |
| 44 | *Medicago laciniata* (L.) Mill | Leguminosae | UOG 349 | + | - | - | + | Mena | BLP | Herb | Whole plant | It is used as diuretic and in kidney diseases. Also used as animal fodder. |
| 45 | *Melilotus indicus* (L.) All. | Leguminosae | UOG 389 | + | - | + | + | Sinjiee | UC | Herb | Whole plant | Externally used as poultice or plaster on swelling and also be used as eye sight tonic. |
| 46 | *Acacia modesta* Wall. | Leguminosae | UOG 355 | + | - | + | + | Phulai | BLP | Tree | Gum | Gum is restorative, as adhesive and used in many food industries. Also used as fodder for grazing animals. |
| 47 | *Acacia nilotica* (L.) Delile | Leguminosae | UOG 356 | + | + | + | + | Keekar | BLP | Tree | Bark, pods and gum. | Astringent, bark used to cure diarrhea. |
| 48 | *Albizia lebbek* (L.) Benth. | Leguminosae | UOG 357 | - | + | + | + | Kala sirin | RT | Tree | Bark | Inflammations, cough, eye infections, Gingivitis, lung problems, pectoral problems, tonic, tumors, hernia and secondary infertility. |
| 49 | *Alhagi maurorum* Medik | Leguminosae | UOG 346 | - | + | - | - | Jawan Janasa booti | RT | Herb | Whole plant | Plants are used as diuretic, expectorant and treatment of freckles and pimples of face. It is also be used as fodder. |
| 50 | *Cassia fistula* L. | Leguminosae | UOG 327 | - | + | - | - | Amaltas/ Golden shower | RT | Tree | Whole Plant | It is used in stomach disorders, common cold, fever and in blood purification. |
| 51 | *Dalbergia sissoo* DC. | Leguminosae | UOG 388 | + | + | + | - | Tali | RMT | Tree | Leaves, root and bark | Stimulant, astringent and alterative. |
| 52 | *Senna occidentalis* (L.) Link | Leguminosae | UOG 328 | + | + | + | + | Desi Sanna | RMT | Herb | Whole plant use | Used to lower inflammation. Leaves are eaten raw to cure piles and also used in respiratory problems and in liver diseases. |
| 53 | *Tephrosia lupinifolia* DC. | Leguminosae | UOG 347 | - | - | + | - | Fish poison | UC | Herb | Leaves and root. | It is used in constipation, inflammation of liver and also against snake poison. |
| 54 | *Trifolium resupinatum* L. | Leguminosae | UOG 350 | - | + | - | - | Jangli shtala | UC | Herb | Whole plant | It is used as vegetable and grazed by cattle for production of milk. |
| 55 | *Bombax Ceiba* L. | Malvaceae | UOG 361 | + | - | + | - | Sumbal | UC | Tree | Whole plant | Treatment of genital organs, gonorrhea, astringent for diarrhea |
| 56 | *Malva parviflora* L. | Malvaceae | UOG 360 | + | + | + | + | Sonchal | BLP | Herb | Herb Leaves and seed | Used to cure common cold, cough and constipation. |
| 57 | *Malva sylvestris* L. | Malvaceae | UOG 359 | + | + | + | + | Khabzi | UC | Herb | Whole plant | Known for cooling, febrifuge and used in urinary bladder problems. |
| 58 | *Melia azedarach* L. | Meliaceae | UOG 358 | - | + | - | - | Dherak | RMT | Tree | Leaves and fruit | Skin infection and skin diseases. |
| 59 | *Morus alba* L. | Moraceae | UOG 366 | + | + | + | - | Sheehtoot | UC | Tree | Fruit and bark | Refrigerant, used for sore throat. bark is purgative. |
| 60 | *Morus nigra* L. | Moraceae | UOG 367 | - | - | + | + | Kala toot | BLP | Tree | Root, leaves and fruit | Bad thorax and stomach worms. |
| 61 | *Broussonetia papyrifera* (L.) L Her. ex Vent. | Moraceae | UOG 362 | + | + | + | + | Jangli toot | UC | Tree | Bark and fruit | Laxative and febrifuge. |
| 62 | *Ficus benghalensis* L. | Moraceae | UOG 363 | - | - | + | - | Borh | UC | Tree | Whole Plant | It is used in the treatment of sexual weakness and in hepatitis. |
| 63 | *Ficus elastic* Roxb. ex Hornem. | Moraceae | UOG 365 | - | - | + | - | Rubber | RT | Tree | Whole Plant | It is used in the treatment of jaundice. |
| 64 | *Ficus religiosa* L. | Moraceae | UOG 364 | - | + | - | - | Peepal | BLP | Tree | Fruit and Seeds | Used against constipation, vomiting, asthma and urinary problems. |
| 65 | *Callistemon viminalis* (Sol.ex Gaertn.) G. Don | Myrtaceae | UOG 371 | + | + | + | + | Bottle brush | BLP | Tree | Whole Plant | It is an antioxidant and rich in flavonoids, containing polyphenols. |
| 66 | *Eucalyptus camaldulensis* Dehnh. | Myrtaceae | UOG 369 | + | + | + | + | Sufeda | BLP | Tree | Leaves | Nose infection, common cold and in throat diseases. |
| 67 | *Eucalyptus globulus* Labill. | Myrtaceae | UOG 308 | + | + | - | - | Chatta Sufeda | UP | Tree | Whole Plant | Influenza, skin tonic, stomach disorders, fuel wood, furniture, fragrance |
| 68 | *Eucalyptus tereticornis* SM. | Myrtaceae | UOG 368 | + | - | + | + | Sufeda | RMT | Tree | Leaves | Common cold and nose infections. |
| 69 | *Psidium guajava* L. | Myrtaceae | UOG 370 | + | - | - | - | Amrood | RT | Tree | Whole plant | Fruits of this plant are used in treatment of stomach disorders whereas timber is used as furniture making and fuel. |
| 70 | *Oxalis corniculata* L. | Oxalidaceae | UOG 372 | + | + | + | + | Khuti booti/ Yell | RT | Herb | Whole plant | Antiscorbic, refrigerant, cooling and stomachic. |
| 71 | *Anagallis arvensis* L. | Primulaceae | UOG 387 | + | - | + | + | Dahber booti | UC | Herb | Whole plant | Used to lowers fever, diuretic and expectorant. |
| 72 | *Fumaria indica*<br>(Hausskn.) | Papaveraceae | UOG<br>351 | + | + | + | + | Papra | UC | Herb | Whole<br>Plant | It is used in pain, nose infection and diarrhea. Plant is also used in dyspepsia, liver diseases and constipation. |
| 73 | *Polygonum avicularia* L. | Polygonaceae | UOG<br>384 | - | + | + | - | Spike weed | RT | Herb | Whole<br>Plant | Treating common cold, fever and also used in cooking. |
| 74 | *Polygonum plebeium* R. Br. | Polygonaceae | UOG<br>383 | - | + | + | + | Cherri hatha, Small knotweed | BLP | Herb | Whole<br>Plant | Plants are used as expectorant, in cold, pneumonia and also used as fodder. |
| 75 | *Rumex crispus* L. | Polygonaceae | UOG<br>386 | + | + | + | - | Kali palak | BLP | Herb | Whole<br>Plant | Used as treating digestive disorders. Also used as vegetable and fodder. |
| 76 | *Rumex dentatus* L. | Polygonaceae | UOG<br>385 | + | + | + | + | Jangli palak | UC | Herb | Whole<br>Plant. | It is diuretic, laxative and consumed as fodder. |
| 77 | *Portulaca oleracea* L. | Portulacaceae | UOG<br>373 | + | - | + | + | Lonak, Kulfa, Purslane | RT | Herb | Whole<br>plant | Plants are used against kidney infection, painful urination, jaundice, iron deficiency and skin allergy. |
| 78 | *Portulaca quadrifida* L. | Portulacaceae | UOG<br>374 | + | - | + | - | Desi Kulfa | UC | Herb | Whole<br>plant | Plants are used in respiratory problems such as asthma, cough and cold. Additionally, as vegetable. |
| 79 | *Ranunculus muricatus* L. | Ranunculaceae | UOG<br>391 | - | - | + | + | Buttercup | RMT | Herb | Whole<br>Plant | Found as ornamental weed and used as fodder. |
| 80 | *Ziziphus jujube* Mill. | Rhamnaceae | UOG<br>390 | + | + | + | - | Ber |  | Tree | Leaves and<br>fruits. | Externally leaves used in boils and scabies. Leaves are astringent. |
| 81 | *Datura innoxia* Mill. | Solanaceae | UOG<br>394 | + | - | + | + | Datura / Thorn Apple | RT | Herb | Seeds | It is used in the treatment of abdominal disorders, poultices, Gonorrhea and as a fodder. |
| 82 | *Solanum nigrum* L. | Solanaceae | UOG<br>395 | + | + | + | - | Kainch Mainch Nights hade | BLP | Herb | Leaves | For abnormal and painful secretions from ears. Also used in blood purification and profuse menses. |
| 83 | *Solanum xanthocarpum* Schrad. & H. Wendl., | Solanaceae | UOG<br>392 | - | + | + | - | Kandiri | UC | Herb | Whole<br>Plant | It is used as antipyretic, abdominal pain and also as fodder. |
| 84 | *Withania somnifera* (L.) Dunal | Solanaceae | UOG 393 | + | + | + | + | Ak San/ Winter Cherry | UC | Herb | Whole plant | Aphrodisiac, alternative. Fruit is diuretic. Tubers used to treat ulcer. It is used in the treatment of reducing cholesterol level and also be used in sexual debility. |
| 85 | *Tamarix aphylla* (L.) H. Karst. | Tamaricaceae | UOG 396 | - | + | + | - | Frash, Okan, Peelish | RMT | Tree | Whole plant | Plant is as tonic and astringent and also be used in skin diseases. Timbers are used in shelters and as fuels. |
| 86 | *Lantana camara* L. | Verbenaceae | UOG 398 | + | + | + | + | Butterfly weed | BLP | Shrub | Stalk & leaves | Plant extract has anti-ulcer effects. Extract from leaves used as antipyretic and carminative. |
| 87 | *Tribulus terrestris* L. | Zygophyllaceae | UOG 399 | + | + | + | + | Bhakra | BLP | Herb | Whole Plant | The herb is used in sexual debility premature ejaculation and also be used in back pain, gonorrhea and urinogenital diseases. |
| Monocotyledons |  |  |  |  |  |  |  |  |  |  |  |  |
| 88 | *Phoenix dactylifera* L. | Arecaceae | UOG 303 | - | + | - | - | Phall wali khajoor | RT | Tree | Leaves, fruit and stem | Seeds are edible and are used as laxative, toothache and also as fuel. |
| 89 | *Phoenix sylvestris* (L.) Roxb. | Arecaceae | UOG 304 | + | + | - | - | Khajoor | BLP | Tree | Leaves, fruit and stem | It is used as laxative, toothache and fuel. |
| 90 | *Cyperus rotundus* L. | Cyperaceae | UOG 337 | - | + | - | - | Deela | RMT | Herb | Rhizomes | Effective in treating fever, diarrhea, and dysentery. Other uses are for blood disorders, indigestion, diarrhea, dysentery, cholera, stomachic. |
| 91 | *Avena sativa* L. | Poaceae | UOG 377 | + | - | - | - | Javi | UC | Herb | Whole Plant | It is diuretic and helps in digestive problems. Also used as fodder and fuel. |
| 92 | *Cynodon dactylon* (L.) Pers. | Poaceae | UOG 382 | + | + | + | + | Khabal | RT | Herb | Whole plant | It is laxative, astringent, and diuretic. |
| 93 | *Desmostachya bipinnata* (L.) | Poaceae | UOG 378 | + | + | + | - | Dabb | RT | Herb | Whole Plant | Uses include treatment of diseases of blood, fodder, fuel and thatching. |
| 94 | *Dichanthium annulatum* (Forssk.) Stapf | Poaceae | UOG 376 | - | - | - | + | Palwan grass | BLP | Herb | Whole Plant | It is used as fodder, fuel, and for thatching purposes. |
| 95 | *Eleusine indica* (L.) Gaertn. | Poaceae | UOG 380 | - | - | + | - | Crow feet grass | RT | Herb | Whole Plant | It is used for blood diseases, laxative, digestive disorders and making baskets. |
| 96 | *Poa annua* L. | Poaceae | UOG<br>379 | - | - | + | - | Chota<br>grass | RMT | Herb | Whole<br>Plant | It is used as laxative, burning sensation,<br>fuel and as fodder. |
| 97 | *Saccharum*<br>*arundinaceum*<br>Retz. | Poaceae | UOG<br>381 | + | + | + | - | Sarkan<br>da | BLP | Herb | Stem<br>and root | Diuretic and diaphoretic, useful in<br>blood troubles. |
| 98 | *Saccharum*<br>*spontaneum* L. | Poaceae | UOG<br>375 | + | + | + | + | Kahi | RT | Herb | Whole<br>plant | Laxative, used in burning sensation,<br>phthisis (pulmonary tuberculosis) and<br>in diseases of blood. |
| 99 | *Typha latifolia*<br>L. | Typhaceae | UOG<br>397 | + | + | + | - | Large<br>marsh | RMT | Herb | Leaves<br>and<br>Pollens | Astringent, sedative and anticoagulant. |
| **Ferns** |  |  |  |  |  |  |  |  |  |  |  |  |
| 100 | *Equisetum*<br>*arvense* L. | Equisetaceae | UOG<br>400 | - | - | + | - | Char<br>booti | UC | Herb | Stem | It is used for the treatment of diabetes. |
| 101 | *Marsilea*<br>*quadrifolia* L. | Marsileaceae | UOG<br>403 | + | + | + | + | Four<br>leaf<br>clove | RMT | Herb | Seeds<br>and<br>leaves | A juice made from the leaves is<br>diuretic, febrifuge and anti-inflammatory and against snake bite. |
| 102 | *Osmunda*<br>*regalis* L. | Osmundaceae | UOG<br>401 | + | + | + | - | Spiral<br>booti | BLP | Herb | Roots<br>and<br>shoots | Used for blood and renal diseases. |
| 103 | *Adiantum*<br>*capillus-veneris*<br>L. | Pteridaceae | UOG<br>402 | - | - | + | - | Maide<br>n hair<br>fern | RT | Herb | leaves | Effective in bronchial disorders. |
| 104 | *Azolla pinnata*<br>R. Br. | Salviniaceae | UOG<br>418 | + | - | + | + | Mosqu<br>ito fern | RMT | Herb | Whole<br>plant | Whole plant is used as a green manure<br>in paddy fields, to add nitrogen and<br>organic matter to the soil, activator of<br>compost making process, and phyto-remediation of environmental<br>pollutants. |
| **Bryophytes** |  |  |  |  |  |  |  |  |  |  |  |  |
| 105 | *Funaria*<br>*hygrometrica*<br>Hedw. | Funariaceae | UOG<br>416 | + | + | + | + | Moss | UC | Folios<br>e | Whole<br>Plant | Effective for cardiovascular diseases<br>and in nervous disorders. |
| 106 | *Marchantia*<br>*polymorpha* L. | Marchantiace<br>ae | UOG<br>415 | + | - | + | - | Umbre<br>lla<br>liverw<br>ort | BLP | Thall<br>oid | Whole<br>Plant | It is used in the treatment of hepatitis<br>and is diuretic. |
| 107 | *Polytricum* sp. | Polytrichaceae | UOG<br>417 | + | - | + | + | Hair<br>moss | RT | Folios<br>e | Whole<br>Plant | It is used for treating bronchial<br>disorders. |
| 108 | *Riccia cavernosa* Hoffm. | Ricciaceae | UOG 418 | + | - | + | + | Spons wat ervorkje | BLP | Thalloid | Whole Plant | It is diuretic and also used in eye diseases. |
| **Aquatic Plants** |  |  |  |  |  |  |  |  |  |  |  |  |
| 109 | *Pistia stratiotes* L. | Araceae | UOG 405 | + | - | + | - | Water lettuce | RT | Weed | Leaves | It is used as antioxidant. |
| 110 | *Spirodela polyrhiza* (L.) | Araceae | UOG 404 | + | + | - | - | Duckweed | RT | Weed | Whole Plant | It is used in the treatment of injury and wound healing. |
| 111 | *Ceratophyllum demersum* L. | Ceratophyllaceae | UOG 406 | + | - | + | - | Coontail Weed | RT | Weed | Leaves | It is used in the regulation of bile secretions and also treatment of scorpion bites. |
| 112 | *Chara Schweinitzii* A. Braun | Charophyceae | UOG 407 | + | + | - | + | Musk grass | BLP | Weed | Stem, Branches | Antiseptic. |
| 113 | *Hydrilla verticillata* (L.F.) Royle | Hydrocharitaceae | UOG 414 | - | + | + | + | Esthwaite waterweed | RMT | Weed | Leaves | It is used in wound healing and in the abscess boil. |
| 114 | *Nelumbium nuciferum* Gaertn | Nymphaeaceae | UOG 409 | + | + | + | - | Indian lotus | UC | Weed | Whole Plant | It is used in cardiovascular and liver diseases and curing pile. |
| 115 | *Nymphaea lotus* L. | Nymphaeaceae | UOG 408 | + | + | - | - | Water lily | RT | Weed | Leaves | Regulates menstruation and cure for digestive disorders. |
| 116 | *Eichhornia crassipes* (Mart.) | Pontederiaceae | UOG 410 | + | + | + | + | Water hyacinth | BLP | Weed | Petiole and leaves | It is antioxidant, carotene rich and used as manure. |
| 117 | *Potamogeton* sp. | Potamogetonaceae | UOG 412 | + | - | - | - | Pond weed | UC | Weed | Leaves and seeds | It is known for treating stomach disorders and having flavoring agents. |
| 118 | *Zannichellia palustris* L. | Potamogetonaceae | UOG 411 | + | - | + | - | Horned pondweed | RT | Weed | Leaves | It is used in the treatment of digestive disorders. |
| 119 | *Spirogyra buchetii* Petit | Zygnemataceae | UOG 413 | - | + | + | + | Green algae | BLP | Weed | whole Plant | Used as a source of antioxidant and mutagenic. |

Vou. No. Voucher Number; RT, River Tavi; UC, Upstream Chenab; RMT, River Munawarwala Tavi; BLP, Bhalolpur. Signs used are as indication of presence (+) or absence (-) of particular species at particular location.

### Ethnopharmacological and industrial resources of the area

Plants including groups, families, botanical names, local names, habit, locality, and plant part used and their application are mentioned in Table 2. Reported flora is being used to support wide range of livelihood activities providing medicine, food, fodder, grazing, timber, fuel and many other services for indigenous community. Although people are getting benefits but in current scenario, members of local community residing in rural areas are preferred to migrate to cities as they are reluctant to live in the rural area, due to modern age facilitates and unexpected flood. Because the flow of water is being controlled by Indian side and they released water as per their will [27]. Whereas on the other hand, in spite of increase in prices for allopathic, they would also like to enjoy herbal plant wealth in the shape of variety of different products such as food, fodder, vegetables, medicines, fibers and aromatic plants, that means it intact with their culture and at any mean few of them wouldn’t like to avoid [28].

#### Application of whole plant and plant parts used as preparatory material

As presented in figure 4, different plant parts are being used in addressing ailments. Whole plant or their parts are being used for various medicinal as well as industrial purposes and they are the integral part of livelihood in the area. Their applications were recorded in herbal medicine, pharmaceutical, food, fodder, grazing, timber and fuel. Further, other provisional services are furniture manufacturing, wooden pots, sports, goods, sheltering, home making and wharfs as wooden bridge, transportation, boats, ships and in railway tracks and so on.

**Figure 4.**
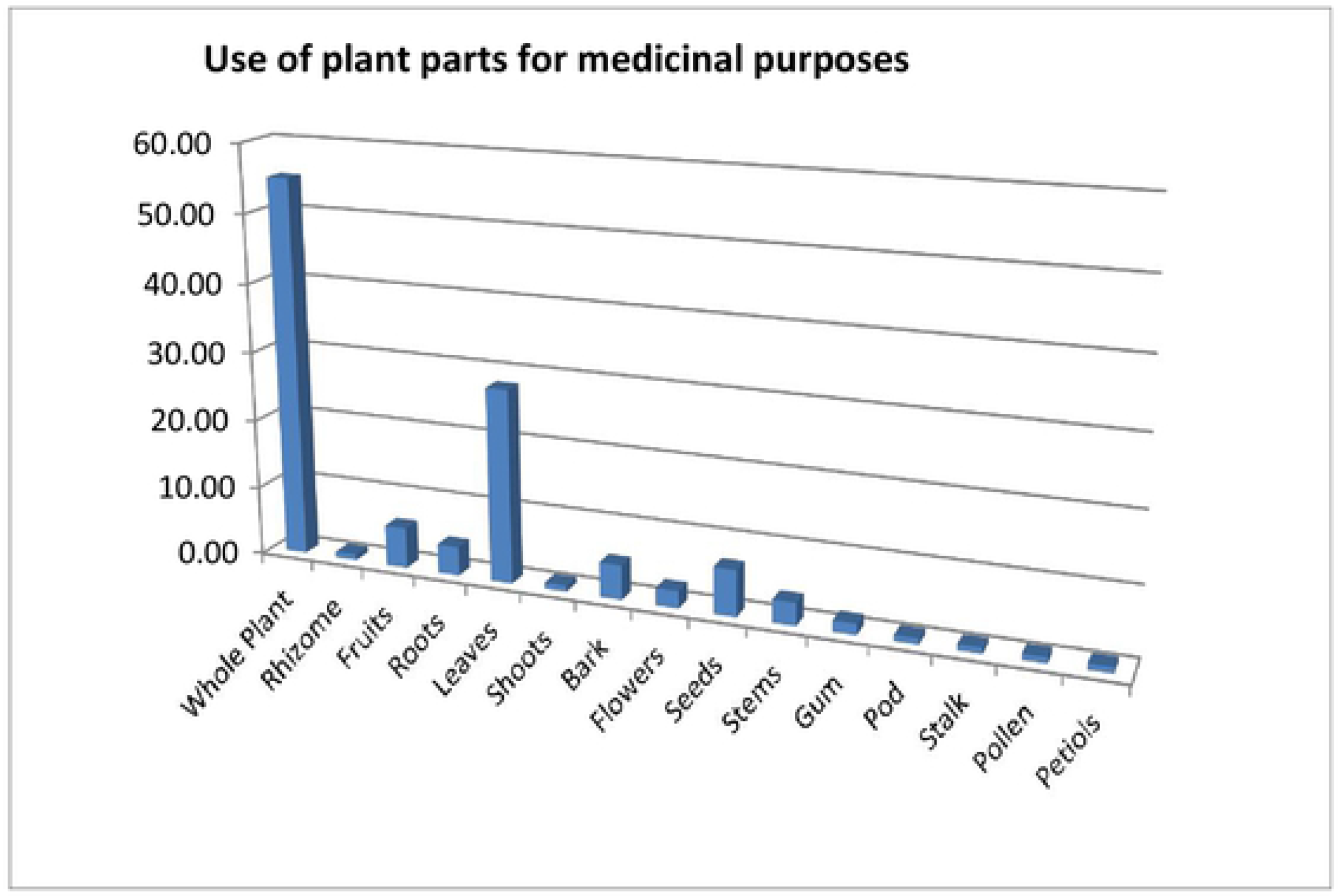
Use of plant parts for medicinal purposes studied at Head Maralla

#### Phytotherapeutic uses and remedies

Across globe, 20% of plant species are being used in health care system [29]. In 19^th^ century, human beings used plants as therapeutic agents and today 80% of the world depends on the medicinal plants for their uses of primary and secondary human health care either directly or indirectly. Since human civilization, 4000 plant species are documented as food while cultivated plants are only 150 of different species. Therapeutically, plant secondary metabolite substances are chemically diverse and main source of drugs and various formulations [30]. Figure 5 showed detail about plants used as digestive, urinary, respiratory, circulatory and general body problems.

**Figure 5.**
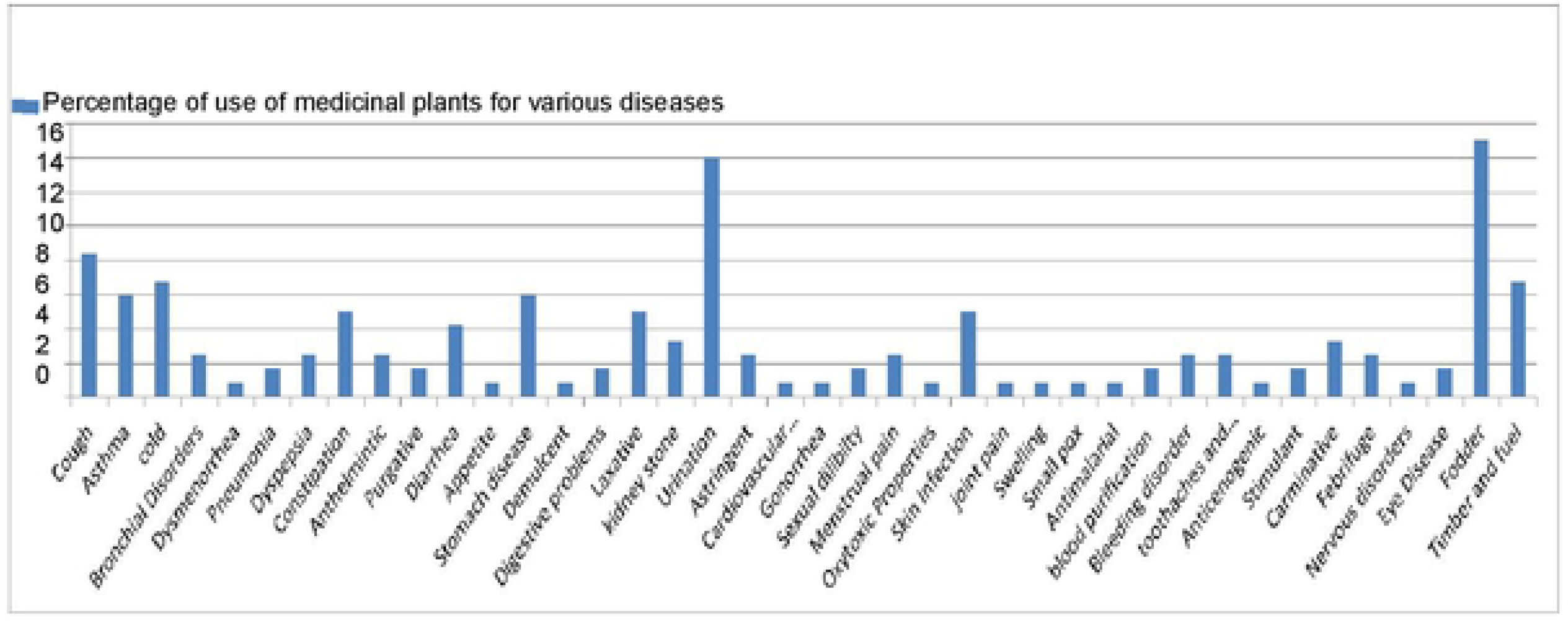
Percentage of use of wild plants for various ailments and industrial uses reported from Head Maralla

### Quantitative analysis of ethnobotanical data

#### Informant Consensus Factor (ICF)

ICF depends on the availability of plants within the study area to treat diseases. The product of the consensus factor is 0 to 1 and further, it showed the categories of ailments and informant consensus factor for each category (Table 3). ICF calculated for 12 ailments viz., respiratory system, throat disease, liver disease, digestive system, constipation, wound healing, kidney disorders, Jaundice, fever, diarrhea, inflammatory problems and laxative. ICF for these 12 ailments were ranging from 0.66 to 0.93 where the 0.66 was calculated for laxative and 0.93 for respiratory system disorders. Arrange ICF was 0.87 which reflects a high consensus among the informants about the use of plants to treat particular ailment. Usually the value of ICF for disease treatment depends on the availability of plant species in that area [31]. The order of ICF was reported for respiratory system (0.93), throat disease (0.92), liver disorders (0.86), digestive system (0.84), constipation (0.83), wound healing (0.83), kidney disorder (0.81), jaundice (0.79), fever (0.75), diarrhea (0.74), inflammatory problems (0.73) and laxative (0.66). Data indicated that respiratory system disease category was the most common in study area, most of the people had knowledge about its cure. They were practicing 10 different species (*Justicia adhatoda* L., *Trianthema portulacastrum* L., *Achyranthes aspera* L., *Calotropis procera* (Aiton) Dryand., *Xanthium strumarium* L., *Senna occidentalis* (L.) Link, *Euphorbia hirta* L., *Leucas aspera* (Willd.) Link, *Ficus religiosa* L., *Portulaca quadrifida* L.).

**Table 3.** Disease category cured through number of taxa/species, reports and informants consensus factor (ICF) for Head Maralla vegetation

| Use categories | Number of taxa/species | Number of use reports | ICF |
| --- | --- | --- | --- |
|  | Nt/s | Nur |  |
| Respiratory system | 3 | 32 | 0.93 |
| Throat diseases | 3 | 29 | 0.92 |
| Liver disorders | 5 | 31 | 0.86 |
| Digestive system | 7 | 34 | 0.84 |
| Constipation | 6 | 31 | 0.83 |
| Wound healing | 6 | 31 | 0.83 |
| Kidney disorders | 6 | 28 | 0.81 |
| Jaundice | 7 | 30 | 0.79 |
| Fever | 8 | 30 | 0.75 |
| Diarrhea | 8 | 28 | 0.74 |
| Inflammatory problems | 9 | 31 | 0.73 |
| Laxative | 8 | 22 | 0.66 |

In addition to respiratory disorders these species were also reported from other areas of Pakistan like Navapind and Shahpur Virkan, district Sheikhupura, Gallies-Abbottabad, Northern Pakistan, Tormik Valley, Karakorum range, Baltistan, Naran Valley, Western Himalaya lying between three different provinces Punjab, Khyber Pakhtun Kkaw (KPK) and Gilgit Baltistan [32–35]. However, two species were used for curing cough and stomach disorders *Adiantum capillus-veneris* L.; decoction of leaves is prescribed in cold, cough, flue and asthma used in the area of New Murree, Patriata, district Rawalpindi and *Cichorium intybus* L., root and leaves were used as appetizer, depurative, digestive, diuretic, laxative and tonic reported from district Bagh, Azad Kashmir[36, 37]. The lowest value of ICF was obtained for laxative (0.66), inflammatory problems (0.73), diarrhea (0.74) and fever (0.79). One reason for low values might be due to lack of communication among informants. Maximum value close to 1 indicated that people were using these species in relatively large quantities, while low value reflected that informants disagree on the specie use against treatment of different ailments.

#### Use values (UVs)

UVs ranged from 0.2 to 6.0, the highest UVs were recorded for *Lepidium didymus* L., (6.0), *Nelumbium nuciferum* Gaertn., (3), *Albizia lebbeck* (L.) Benth. (2.4) and *Cyperus rotundus* L., (2.25), *Amaranthus viridis* L. (2), *Ficus religiosa* (2), *Ficus benghalensis* L., (2), *Calotropis procera* (2), *Lepidium didymus* (6), *Cleome viscose* L., (2), *Tinantia erecta* (Jacq.) Fenzl (2), *Cucumis melo* L., (2), *Alhagi maurorum* Medik., (2), *Mentha spicata* L., (2), *Marchantia polymorpha* (2), *Riccia cavernosa* (2), *Hydrilla verticillata* (L. F.) Royle (2) and *Potamogeton* sp., (2), respectively (Table 4). Plants with high UVs were also reported to be used in various parts of Pakistan, these plants have potential to develop herbal drugs along/after proper phytochemical and pharmacological screening. Focus on plant species with high UVs advocated plant resource sustainability and conservation. Lowest UVs were reported for *Acacia modesta* Wall., (0.2), *Chrozophora tinctoria* (L.) A., Juss., (0.3) *Polytricum* sp. (0.3), *Pistia stratides* (0.3) and *Alternanthera pungens* Kunth (0.3) [38]. It was noted that number of informants was not familiar with these plant species as far as their ethnobotanical uses are concerned. Lower UVs indicated existence of poor knowledge about particular species in the study area [40, 41].

**Table 4.**
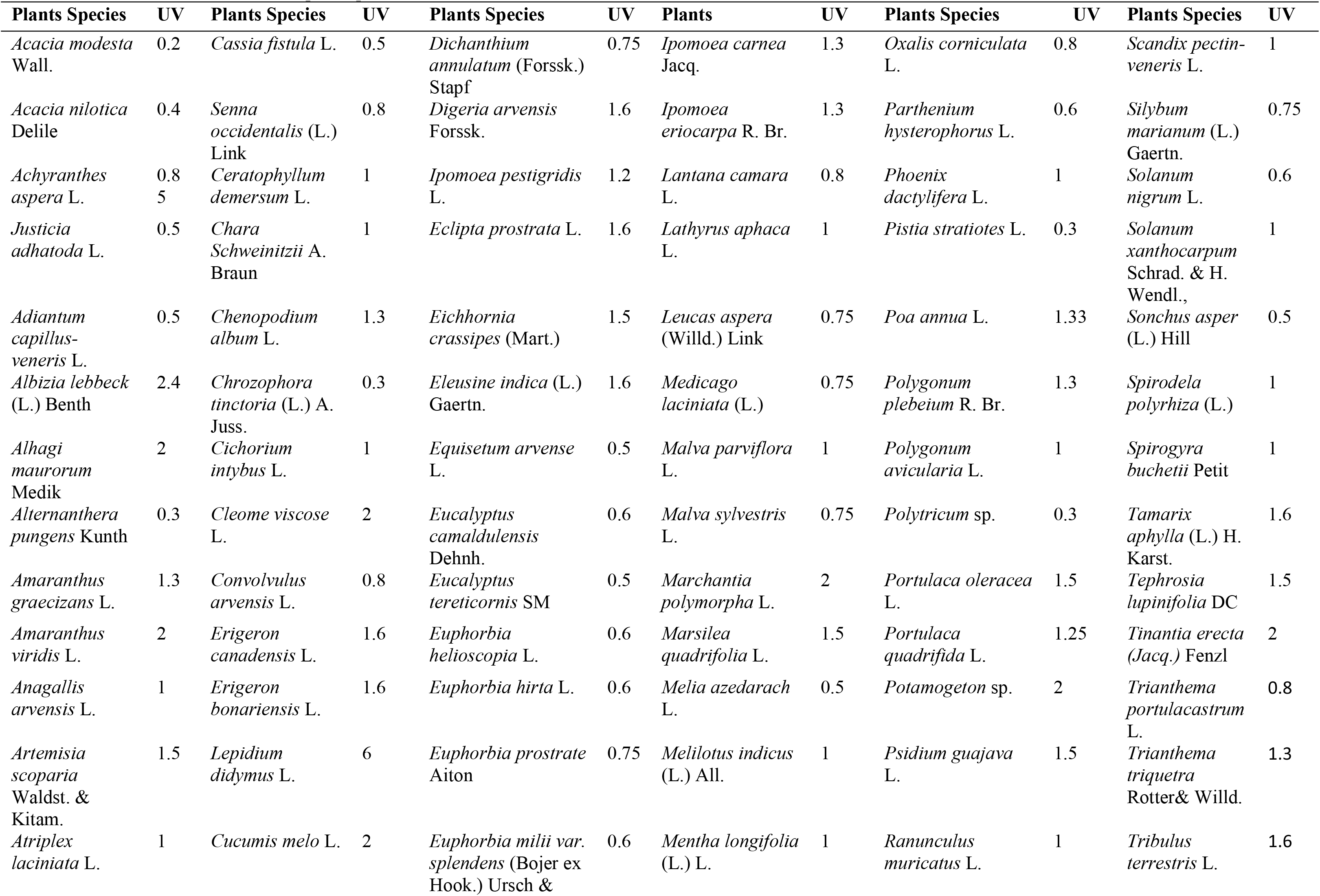
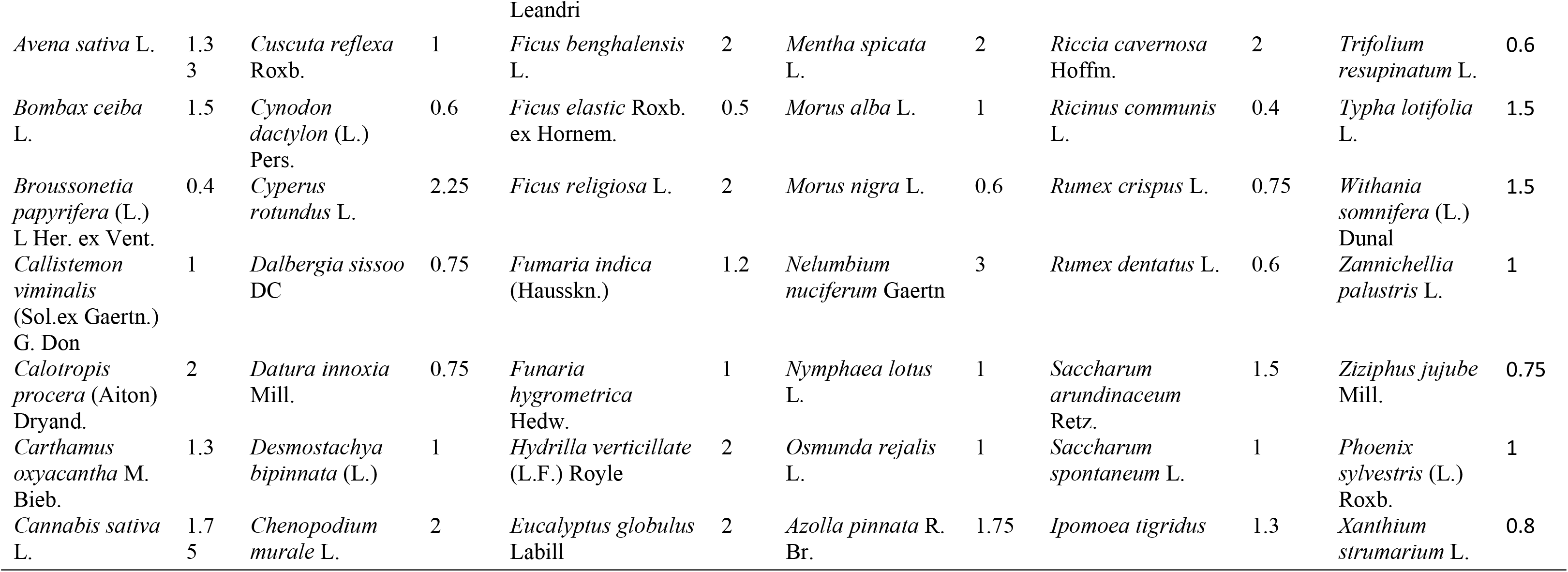
Various plant species with their Used Values (UV) recorded from Head Maralla

#### Comparison with neighboring areas

For comparison ethnopharmacological & ethnobotanical data from 40 published studies of adjacent areas were consulted of which few of them were found with related/same species (Table 5). Many species were found with new applications. Ethnobotanical studies of surrounding districts, Central, Southern and Northern Punjab, Sindh, KPK, Baluchistan, Azad Kashmir, Iran, India and Bangladesh were compared with Head Maralla vegetation. Data from some African countries were also compared having similar type of geoclimatic and topographic conditions. Few species were common in many reports that reflects cross cultural knowledge or frequent exchange of material beyond boundaries in past. Consequently, this exchange of knowledge afterwards converted into traditional medicinal system and was transferred from generation to generation.

**Table 5.** Comparison with adjacent regions national as well international reports published

| S. No. | Area of studies | Study Year | Number of recorded plant species | Families | Total species common in both areas | Plant with similar uses (%) | Plants with dissimilar uses (%) | Species enlisted only in study area | Jaccard Index | Citations |
| --- | --- | --- | --- | --- | --- | --- | --- | --- | --- | --- |
| 1 | Wana, South Waziristan, Pakistan | 2013 | 50 | 30 | 16 | 32 | 68 | 102 | 11.76 | Ullah et al., 2013 |
| 2 | Samhani valley Azad Kashmir, Pakistan | 2009 | 120 | 46 | 34 | 27.5 | 72.5 | 84 | 20 | Hussain and Chaudhary, 2009 |
| 3 | Coastal areas of Pakistan | 2014 | 54 | 27 | 19 | 34.6 | 65.4 | 99 | 14.17 | Qasim et al., 2014 |
| 4 | Nara Desert, Sindh, Pakistan | 2008 | 6 | 1 | 3 | 50 | 50 | 115 | 2.54 | Qureshi and Bhatti, 2008 |
| 5 | Kotli AJK, Pakistan | 2014 | 33 | 25 | 2 | 6 | 94 | 116 | 1.36 | Amjad and Arshad, 2014 |
| 6 | Khunjrab National Park Gilgit, Pakistan | 2011 | 43 | 28 | 6 | 13.95 | 86 | 112 | 4.34 | Qureshi et al., 2011 |
| 7 | Khushab, Pakistan | 2011 | 48 | 32 | 26 | 54.5 | 45.5 | 92 | 22.8 | Qureshi et al., 2011 |
| 8 | Hingol National Park Baluchistan, Pakistan | 2012 | 39 | 22 | 10 | 25 | 75 | 102 | 7.63 | Qureshi, 2012 |
| 9 | New Murree, Patriata, District Rawalpindi | 2013 | 93 | 56 | 28 | 30 | 70 | 90 | 18 | Ahmed et al., 2013 |
| 10 | Karak, KPK | 2014 | 160 | 50 | 51 | 31.8 | 68.2 | 67 | 28.97 | Khan et al., 2014 |
| 11 | Tehsil Kabal, Swat, KPK | 2015 | 45 | 27 | 19 | 42.2 | 57.8 | 99 | 15.2 | Khan et al., 2015 |
| 12 | Tehsli Chakwal | 2009 | 29 | 18 | 21 | 72.4 | 27.6 | 97 | 20 | Qureshi et al., 2009 |
| 13 | Tehsil Kharian , District, Gujrat | 2014 | 50 | 32 | 25 | 51.5 | 49.5 | 93 | 21.18 | Ajaib et al., 2014 |
| 14 | Ladha Sub Division, South Waziristan agency, Pakistan | 2016 | 82 | 42 | 25 | 30.4 | 69.6 | 93 | 16.66 | Aziz et al., 2016 |
| 15 | Bhimber, Azad<br>Jamu Kashmir | 2011 | 38 | 22 | 23 | 60.5 | 39.5 | 95 | 20.9 | Mahmood et al., 2011 |
| 16 | Himalayas<br>Mountain Naran<br>Valley | 2013 | 101 | 52 | 6 | 5.94 | 94.16 | 112 | 2.89 | Khan et al., 2013 |
| 17 | District Mirpur<br>AJK, Pakistan | 2011 | 29 | 20 | 16 | 55 | 45 | 102 | 13.91 | Mahmood et al., 2011 |
| 18 | District Gujrat | 2013 | 22 | 18 | 7 | 31.8 | 68.2 | 111 | 5.5 | Parvaiz, 2014 |
| 19 | District Sialkot | 2014 | 18 | 13 | 5 | 27.7 | 72.3 | 113 | 3.96 | Ikram et al., 2014 |
| 20 | Valley Alladand<br>Dehri, Tehsil<br>Baathkhela,<br>District Malakand | 2013 | 92 | 53 | 24 | 26 | 74 | 94 | 14.81 | Alamgeer et al., 2013 |
| 21 | District Swabi,<br>Pakistan | 2017 | 66 | 41 | 15 | 22.7 | 77.3 | 103 | 10.13 | Rozina et al., 2017 |
| 22 | District Kotli,<br>Azad Jamu<br>Kashmir, Pakistan | 2010 | 38 | 25 | 5 | 13.1 | 86.9 | 113 | 3.42 | Ajaib et al., 2010 |
| 23 | Mianwali District | 2007 | 21 | 16 | 13 | 61.9 | 38.1 | 105 | 11.5 | Qureshi et al., 2007 |
| 24 | Tehsil Dargai,<br>District Malakand | 2013 | 40 | 26 | 9 | 22.5 | 77.5 | 109 | 6.42 | Zaman et al., 2013 |
| 25 | Sudhan Gali and<br>Ganga Chotti<br>Hills, District<br>Bagh, Azad<br>Kashmir | 2007 | 33 | 17 | 1 | 3.03 | 96.6 | 117 | 0.67 | Qureshi et al., 2007 |
| 26 | Hattar Region,<br>District Haripur,<br>NWFP | 2008 | 45 | 26 | 29 | 64.4 | 35.6 | 89 | 27.61 | Hussain et al., 2008 |
| 27 | Tehsil Kotli<br>Sattian, Soan<br>River, Rawalpindi | 2016 | 35 | 20 | 13 | 37.14 | 62.85 | 105 | 10.23 | Bashir and Ahmad, 2016 |
| 28 | District Kotli<br>Azad Jamu<br>Kashmir | 2014 | 93 | 46 | 26 | 27.96 | 72.05 | 92 | 16.35 | Ajaib et al., 2014 |
| 29 | Neelum Valley,<br>AJK, Pakistan | 2011 | 40 | 31 | 5 | 12.5 | 87.5 | 113 | 3.37 | Mahmood et al., 2011 |
| 30 | Eight Districts of<br>Punjab, Vehari, | 2012 | 48 | 23 | 16 | 33.33 | 66.66 | 102 | 11.94 | Zereen and Khan, 2012 |
|  | Pak Pattan,<br>Lahore,<br>Faisalabad,<br>Nankana Sahib,<br>Sahiwal, Sialkot,<br>Narowal |  |  |  |  |  |  |  |  |  |
| 31 | Vehari, Pak<br>Pattan, Lahore,<br>Faisalabad,<br>Nankana Sahib,<br>Sahiwal, Sialkot,<br>Narowal and<br>Central Punjab | 2013 | 51 | 5 | 7 | 13.72 | 86.27 | 111 | 4.51 | Zereen et al., 2013 |
| 32 | Field survey of<br>the Punjab | 2012 | 36 | 13 | 5 | 13.88 | 86.11 | 113 | 3.47 | Madani et al., 2017 |
| 33 | Erer valley of<br>Babile Wereda<br>Eastern Ethiopia | 2012 | 51 | 28 | 4 | 7.84 | 92.15 | 114 | 2.48 | Belayneh et al., 2012 |
| 34 | D.I. Khan, Bannu,<br>Lakki Marwat,<br>Karak and Kohat<br>KPK | 2015 | 52 | 36 | 19 | 36.5 | 63.4 | 99 | 14.93 | Tariq et al., 2015 |
| 35 | Indus River, dera<br>Ismail Khan,<br>Pakistan | 2014 | 70 | 39 | 26 | 37.14 | 62.85 | 92 | 19.11 | Mussarat et al., 2014 |
| 36 | Western Uganda | 2014 | 231 | 72 | 2 | 0.86 | 99.13 | 116 | 0.57 | Asiimwea et al., 2014 |
| 37 | Manila<br>Philippines | 2017 | 76 | 40 | 5 | 6.57 | 93.42 | 113 | 2.71 | Balinado and Chan,<br>2017 |
| 38 | Susunia hill of<br>Bankura District,<br>West Bengal,<br>India | 2015 | 25 | 17 | 5 | 20 | 80 | 113 | 3.75 | Rahaman and<br>Karmakar, 2015 |
| 39 | Kalenga Forest<br>Area Bangladesh | 2014 | 35 | 25 | 6 | 17.1 | 82.85 | 112 | 4.25 | Uddin and Hassan, 2014 |
| 40 | Samburu<br>Community<br>Wamba, Kenya | 2014 | 33 | 5 | 1 | 3 | 97 | 117 | 0.67 | Omwenga et al., 2014 |
| Average |  |  | 60 | 29 | 14 | 29.06 | 70.91 | 103 | 11.3 |  |

## 2. Discussion

### Diversified flora of Head Maralla

Flora of Head Maralla is diversified due to deposition of minerals and soil by flood water. The present study is the first comprehensive and systematic ethnobotanical survey of Head Maralla covering crosslinks of Gujrat and Sialkot districts on one side and Indian Occupied Kashmir and Jammu on the other side. Findings were also compared with available reports in adjacent districts along with some of neighboring countries. Various environmental factors affected vegetation like soil characteristics and flood water [43, 44]. Plants were distributed at all sites and usually available in particular season. During summer long day plants are growing. In monsoon, rainfall ratio is highest in Sialkot, as compared to rest of the country and it promotes vegetation. In winter short day plants are growing that require sunlight in small quantity for example *Ranunculus muricatus* L., *Eclipta prostrata* (L.) L.*, Anagallis* arvensis L., and *Oxalis corniculata* L.

#### Impact of wild plants on local community health and well being

Reported wild plants have potential to treat various life threatening diseases like ulcer, tumor, diarrhea and asthma, etc. Likewise use of wild plants as medicinal and aromatic purposes are also documented from various areas of the country, need is to explore the hidden potential on scientific basis. Owing to that, concentrated efforts would be required and community mobilization toward conserving local wild plant resources [45]. Studies related to Mardan area showed plants such as *Acacia nilotica* Delile, *Morus alba* L., *Luffa cylindrical* L. (Rox), and *Fagonia cretica* L., were reported based on phytochemical analysis and their uses as medicinal, herbal and functional foods [46].

Following plant species were also reported *Calotropis procera*, *Melia Azedarach* L., *Azadirachta indica* A. Juss, *Artemsia* sp., *Neolitsea chinensis* (Gamble) Chun, *Eucalyptus* sp., *Punica granatum* L., and *Nigella sativa* L. These species are attributed curative effects against various maladies including antimicrobial activities. These plants were also documented from various regions of Pakistan (Khyber Pakhtun Khaw, North of Punjab, and Azad Jammu and Kashmir) [47]. It is evident that screening of potential antimicrobial agents is prerequisite for evaluating therapeutic potential leading isolation of new bioactive compounds [48]. A study based on ethnobotanical data of Morgah Biodiversity Park, Rawalpindi, Pakistan, was conducted on 40 species belonging to 39 genera of 32 families also showed local inhabitants are benefiting themselves [49]. Furthermore, lifestyle changes now modify living by the availability of alternatives to local herbs along, it declines herbal uses. Numbers of synthetic materials are available in market as substitute to natural products. Likewise, furniture was earlier made up of wood obtained preferably from *Delbergia sessio* DC., now replaced by synthetic material like plastic, iron, plywood etc., the demand for such type of furniture and households reduces now is not only due to shortage of raw material but also of high cost. While interestingly still today, Gujrat and Chiniot districts are famous for furniture manufacturing in Pakistan and attracts markets of Europe. Windows and doors made of *Cedrus deodara* (Roxb.) G. Don., are also very popular but cost raises many folds. Therefore, young generation is shifted to use synthetic material. Another picture is Sialkot which is very famous for manufacturing of sports goods and musical instruments but now shifted to alternative material. One solid reason is cutting of wood trees and reduction in forests’ timber and lumber lack of capacity to support local industry. So, threats to loosen traditional knowledge and skill labor are also at the verge of extinction. Such paradigm shift complies with documenting ethnobotanical knowledge of the concerned area. It is therefore important to document the ethnobotanical data before the information is lost.

### Informants Consensus Factor (ICF)

Respiratory system disorders and throat diseases were prevalent in the study area which could be attributed to limited availability of hygienic food and drinking water [50–51]. Plants are being used frequently to treat such disorders based on fact that active ingredients are available in these plants. Various ethnic communities across globe are employing traditional knowledge to treat respiratory and throat disorders [32, 34, 35]. Ethno pharmacological studies have shown that in some parts of the world, respiratory system disorders and throat diseases are come under a ß category[32, 33]. Although, Head Maralla have never been studied ethnobotanically and ethnomedicinally, current findings little bit agrees with previous reports related to adjacent areas [32, 52], while particularly supporting the results of Kayani et al, 2014 [34], that respiratory disorders and throat infections were dominant diseases in Gallies of Abbottabad, a city in Himalyan range of Northern Pakistan.

High ICF values obtained in this study indicated a reasonable high reliability of informants on the use of medicinal plants particularly for respiratory system disorder and throat diseases complaints, while low ICF values for laxative indicated that informants have little knowledge [53]. Frequently a high ICF vale is allied with a few specific plants with high use reports for treating a single disease category while low values are associated with many plant species with almost equal or high use reports, suggesting a low level of agreement among informants or the use of reported plant species to treat a particular disease category [54].

#### Use values (UVs)

In present studies, *Lepidium didymus*, *Nelumbium nuciferum* Gaertn., *Albizzia lebbek*, *Cyprus rotundus* were found with high UV. It is found that plants having high use reports (UR) always have high UVs while those plants having fewer URs, reported by informants carry lower UV. It is also observed that plants that are used in some repetitive manner are more likely to be biologically active [55].

The values for UV are important and change with the change of an area and particularly on the basis of knowledge intact with people, therefore values of UV may varies from area to area and even within the same area. On contrary, plants with lower UV value are not necessarily unimportant, which may provide an indication that the young people of the area are not well aware about the application of these plants. Based upon this understanding, it can be assumed that uses of plants and their relevant knowledge is at risk of not being transmitted to further generations, thus this knowledge may eventually disappear [56]. However, it is one view but the authors do not agree with such findings. With the passage of time, more and more exploration of natural compounds and their evaluation from different plant species, it is ascertaining that the role of traditional knowledge is more prevalent and important. At the same time one system of medication may fail and other is luckily available to fill the gap and provide alternative remedy. Another advantage is now ethnobotanical knowledge is scientifically evaluated, validated and further scientific approaches are being used to get authentic results and pure compounds to make new drugs more useful and improving old recipes to cater more productive outcome. It is not the matter of young generation but of health in the community, society structure of Pakistan, where per capita income is still far behind developed countries and people of all creeds consult herbalists, hakims, doctors, religious scholars for all possible solution and improved health care. Additionally, government is also paying attention for improvement of health and health services, therefore devising laws and regulations not only for medical profession but also for other professions that are at par.

This is the first quantitative ethnobotanical investigation to be carried out in the study area, therefore we compared our results with similar quantitative studies carried out in other parts of the country [57–59]. This revealed that there were differences in most of the cited species and their quantitative values. In a study carried out by Abbasi et al [40], *Ficus carica* and *Ficus patrata* were the most cited species, while Bano et al, (2014b) reported that *Hippophae rhmnoides* had the highest use value (1.64) following by *Rosa brunonii* (1.47) [59]. These differences are mostly accounted for high variation in vegetative and geo-climatic conditions of the area and emphasis would be to explore more adjacent areas and direct interaction with local communities.

#### Comparison with other studies conducted in adjacent areas

Few scholars compose formulations and recipes but are very limited and restricted to few plant species. Likewise, wild plant resources and traditional knowledge spread in a very unique way, particular class of people were involved in this business, village man collected plants as wild species and knowledge transfer restricted to only to their own heir not general public. No sanitary or phytosanitary laws were existed in past, even today laws are not followed properly. Hence last few decades tinted that lack of interest in youth created gap in knowledge transformation too, that ultimately increases significance to document immediately available traditional knowledge about wild plants and their applications. The first document related to 1500 medicinal plant species was made from West Pakistan [60, 61].

It is noted that different plant species are being used for treatment of various types of ailments. At a time one plant specie can be used alone or in combination of plants or blend of different compounds used to treat different ailments. Therefore, various plant recipes were used to cure ailments in typical areas. Interestingly it was also observed that some of the plants have specific application while others have multiple uses like *Cassia fistula* L. and *Tamarindus indica*. One tea spoon amaltas fruit pulp and an equal amount of tamarind in one cup of water, left overnight mashed and strained is used for stomach problem. Similarly, one plant specie can be used against more than one disease like *Amaranthus spinosus* is used as antidote and as well as for constipation. Moreover, a single ailment can be treated with a list of plant species like asthma. Many wild plant species possess medicinal as well as industrial applications. *Eucalyptus globulus* Labill. is used both as ingredient in food items, part of medicinal formulations and also for manufacturing furniture and sport goods. Results of present study revealed that *Saccharum spontaneum* L., of family poaceae was one of the specie explored for the first time for various medicinal uses, diseases such as laxative, burning sensation, phthisis (pulmonary tuberculosis) and in diseases of blood. This specie has not been reported before such medicinal properties from any region of the Pakistan, India, Nepal, Iran, Morocco, Singapore and Kenya [62–69]. Eight species belonged to family leguminosae were recorded from different sites of Head Maralla. This family is well known for pharmaceutical and food purposes for both man and livestock. Members of this family are rich in phytochemicals like tannins, phenol, alkaloids, conmarins, glycosides, lignin, quinones and steroids. Along with ethnomedicinal properties it is also a source of protein, timber, fiber, gums, resins, coloring matters, insecticides and molluscicides [70, 71]. Fodder species of this family were also employed for treatment of livestock during fever, stomach diseases, blood purification and cough, respectively. Current findings might lead to the discovery of new drug. Moreover, they might also use for exploration of active phytochemicals to give alternative to compensate the drug requirement. Previous studies conducted in Khunjerab National Park, Gilgit Pakistan, and Ladha subdivision, South Waziristan Agency, Pakistan reported that the members of this family are being used for milk increase in livestock, skin infection, antispasmodic and expectorant. They have estrogenic effect helping in control of menopausal complaints [72, 73].

Three species *Malva sylvestris* L., (Khabzi), *Malva parviflora* L., (Sonchal) and *Bombax ceiba* L. (Sumbal) belong to family Malvaceae. This family is characterized by large number of phytochemicals including tannins, polysaccharides, coumarins, flavonoids, malvin, folic acid and terpenoids, vitamin A, vitmin C, and vitamin E [74]. These phytochemicals made them suitable for treating various human and animal ailments. These three species are being used to treat cooling, febrifuge, urinary bladder problems, common cold, cough, constipation, problems of genital organs, gonorrhea, astringent, and diarrhea etc. Zereen and Khan, [75], while studied important trees of central Punjab, mentioned *Tecomella undulata* (Roxb.) Seeman. Tree bark possesses medicinal properties, this is a new finding from current study [75]. There is needed to further investigate it for active constituents of pharmaceutical significance. Likewise species *Artemisia scoparia* Waldst. & Kitam., *Xanthium stromarium*, *Eclipta alba*, *Eclipta prostrata*, *Carthemus oxycantha* (family compositae) known to be used against skin diseases, as fodder, ornamental, stomach disorders, demulcent, in small pox, malaria, tuberculosis, kidney diseases, allergy, athlets’s foot and ringworm. Further, bleeding disorders, jaundice, fever, toothaches, ulcer and also air tonic. This family is rich in many active compounds like saponin, glycosides, steroids, tannins, diterpenoids, triterpenoids and flavonoids [76]. This means subject family members are being used against ailments of humans and animals as well. Only one study reported the uses of *Carthamus oxyacantha* M. Bieb., as its leaves in small doses are being used for curing intermittent fever, against wounds, ulcers and chornic sores, poultice of slightly roasted leaves administered to reduce inflamed swellings and rheumatism. Fermented leaves relieves chest pain and cure tympanitis whereas current studies found new applications like seed oil is being used for addressing ulcer, toothache, and itch. Therefore, new sources of medication could be explored for preparing new drugs singly or in combination [77].

Commonly grown species like *Mentha spicata* (podina), *Mentha longifolia* (L.) and *Leucas aspera* belong to family lamiaceae used as stimulant, carminative, anti-inflammatory, in digestive/gastric disorders, asthma, jaundice, chronic diseases and in addition to that as salad and fodder. The family is well known to contain various alkaloids, tannins, flavonoids, terpenoids and carbohydrates [78]. These species were reported to be used since centuries and a part of local culture. Previous studies found that fresh leaves are edible, and could be used as carminative, against diarrhea, dysentery and colics [79]. Whereas Current studies find first report from Head Maralla and new applications of *Leucas aspera* (Willd.) Link, which showed that it was being used against asthma, jaundice and chronic diseases. Present investigation provides baseline information to screen out biological activities of these valuable plants in order to develop new antiseptic and insecticidal medicines. At Head Maralla *Chrozophora tinctoria* (family Euphorbiaceae) was being used for wound healing while in literature, it was reported for jaundice, typhoid, seeds cause vomiting and plant yields coloring matter [80]. In another study it was being used as stomachic for chest burning [81]. Present study indicated that *Artemisia scoparia* is used for skin diseases, as fodder and ornamental, whereas it is found diuretic in nature [32], however it has been reported for earache, cardiac problem, fever and blood pressure [71, 82, 83]. People were using *Erigeron bonariensis* L. for treatment of diarrhea, bleeding and hemorrhoids. Further leaves were used treating dysentery and diabetes. While it is reported to be used against asthma and ulcer in the area of Navapind and Shahpur Virkan [32]. In literature it is reported for hemorrhage and as diuretic [6, 71, 84]. *E. prostrata* was discovered as treating allergy, athlete’s foot and ringworm. It is also used in blood purification and urticarial. In literature it is found to be used as antidote of scorpion, and as laxative [32, 85]. *Parthenium hysterophorus* L. is used for skin disorder and often taken internally as a remedy for wide range of aliments. While in previous studies it is found that it can be used as blood purifier, backache, dysentery, fever, and toothache [32, 84, 86, 87]. Another plant specie *S. marianum* was used in liver disorders, chronic inflammatory and in heart burn complaints. Moreover it is found effective as anticancer, wound healing and for tuberculosis [32, 41]. *E. helioscopia* member of family Euphorbiaceae expected to have large number of alkaloids was used for eruption and cholera treatment but in the study Shahpur Virkan and Navapind, it is reported as used for cancer and cholera treatment [32].

*Ficus religiosa* was locally used against constipation, vomiting, asthma, and urinary problems, as in literature it was reported as laxative, and wound healing, diarrhea, ulcer, molar pain and cardiac problems [32, 88, 89]. *Ficus benghalensis* is reported to be used as in the treatment of sexual weakness and in hepatitis whereas previous literature mentioned that it is an important rubber yielding specie could be used for diarrhea, blood purifier and for diabetes [32, 90, 91].

Current investigation found another important plant *Portulaca oleracea* L. (Green purslane), local people used this against kidney infection, painful urination, jaundice, iron deficiency and skin allergy. While previous studies revealed it as valuable in treating urinary and digestive problems only. The diuretic effect of juice makes it useful in elimination of bladder ailments. Plant mucilaginous properties also made it soothing remedy for gastrointestinal problems such as dysentery and diarrhea. Seeds are demulcent, diuretic and vormifuge, these reports are advocating current findings [92]. It was found that *Digeria arvensis* Forssk., was used in headache, burning, dyspepsia, urination and in constipation. Moreover, it is used as salad, vegetable and as fodder. Whereas the foliage of the *Digeria arvensis* was used as potherb for treating constipation at Khushab areas[93]. It is interesting to note that the area of Head Maralla is climatically different from Khushab that is more or less arid and semi-arid area, therefore the local people used these plants for managing different needs. New findings of utilization can certainly open new doors for phytochemists, pharmacologist and even local herbalists to develop new medicines and get benefit from the nature’s treasure of health. Likewise *Cucumis melo* reported from Hingol National Park Baluchistan, Pakistan revealed that ripened fruit is eaten and reported as mild laxative, used in constipation [94]. On the other hand current studies find that *C. melo* was used in tonic and as fodder for livestock in the area of Head Maralla. This report may lead toward discovery of new fodder source.

Published data revealed *Adiantum capillus-veneris* is valued for various ailments like decoction of leaves, also prescribed in cold, cough, flue and asthma [36]. Whereas current studies unveiled that it was used for bronchial disorder in the region of Head Maralla. It is also a discovery of new application of this plant specie. Additionally, the pharmacological properties of *Convolvulus arvensis* L. that represented the uses in Head Maralla region as plant is being used in constipation, headache, wound healing and in menstrual bleeding are new discoveries. In contrary, people from high mountainous ranges of Northern Himalayas of Naran Valley reported that *Convolvulus arvensis* L. powder is considered as purgative & used in evacuation of bowels [35]. Further investigation is suggested for this herb to explore new phytochemicals leading development of new drugs. Earlier reports from Sialkot district revealed that *Eichhornia crassipes* (Mart.), was used in skin treatment and against goiter problem but current study from Head Maralla explored it as multipurpose plant as antioxidant, carotene rich and as manure [95]. Therefore, its traditional knowledge should be evaluated from not only pharmaceutical point of view but also as source of supplementary food and natural manure for vegetables and crop plants are suggested.

#### Recipes used in Head Maralla area

There were number of recipes used by local people for their day to day needs. Crushed leaves of *Mentha spicata* L. and their extract are recommended to use two to three times orally for removing stomach and digestive disorders. Crushed leaves are also used as ‘chatni’ for eating purposes, keeping digestion efficient, and aroma that developed from small pieces of the *mentha*. Chatni is used as coolent and digestive which also included pieces of unripend mango fruits to develop taste. Leaves of *Mentha longifolia* L. are boiled in water and used as effective in reducing mouth smell. Crush fruit/seeds of *Melia azedarach* (Dherak) were also being used against bowls and to remove swelling. Leaves paste is also used against skin disorders. Another common recipe of this plant is as, whole leaves boil for half an hour and then rape in piece of cloth, may applied on wound for healing and removing effects of insect bites. In some cases crushed seed extract was also used as tonic against pimples and other long term skin infections and diseases. *Chenopodium album* L. (bathu) is being used to make vegetable known as ‘saag’, just like *Brassica campestris*. Crushed leaves are mixed with palak (Spinch) and add spices and then boiled for one and half hour to make paste like vegetable, saag. The taste can be enhanced with the use of butter. Another recipe was of *Justicia adhatoda* L., according to traditional knowledge, take about ten leaves and then boil them in water for about half an hour. Juice may be obtained from mixture and then mix one teaspoon of honey into it. It is recommended to take one teaspoon for three times in a day. It will provide relief against cough. Its root ash along with honey can be used as best remedy against asthma. Best results could be obtained while using it at sleeping time.

Another wild plant *Achyranthes aspera* belongs to family Amarantheceae can be effectively used against joint inflammation by making paste of roots and lap for four to five hours. Root paste and boiled extract can also be used against itching, skin infections. The leaves and flowers of the *Scandix pectin-veneris* L. were used as salad and vegetable. Small pieces of leaves were also used as flavoring on different dishes. Further, extract of *Eclipta prostrata* leaves were used as hair tonic which give shiny, dark and healthy hairs. It also helps in reducing baldness and rejuvenates hairs. This discovery could lead to develop new cosmetic products. Another important plant was *Chenopodium murale* (Krund), as whole plant can be used as insect/pest repellent. Usually few bunches of leaves can be place in grain storage places. The latex of *Euphorbia helioscopia* L., (gundi buti) was applied to eruption while seeds with pepper were given in cholera. Moreover, paste of *Euphorbia prostrata* Aiton, in combination with lemon extract was being used to cure skin diseases, itching and for ringworms on the body. *Alhagi maurorum* plants were employed for treatment of freckles and pimples on the face. Pieces of branches (small sticks) are being used for cleaning of teeth or as substitute of tooth brush and tooth paste and are known as ‘Muswak’. Six to seven seeds of *Albizia lebbeck* (L.) Benth (Kala Shirin) were used to eat orally and proved effective against planters. Seeds can take at the interval of one day for a week.

Another specie was *Ipomoea carnea* Jacq., which was used in Jaundice, according to literature. Its seed mixed with sugar and castor oil to treat intestinal pain and worms. Seeds mixed with vinegar were used against swelling [7]. Current findings revealed the anti-carcinogenic and oxytoxic properties, remedy for asthma, bug bites and burns. Whole plant extract, roots, leaves and seeds have biologically active ingredients like Alkaloids, phenolic compounds and glycolipids that have strong impact when applied separately or in a mixture against poisonous insect bite, dog bites, in boils and carbuncles. Poultice of leaves could be used to cure boils, sore and pimples.

*Cichorium intybus* L. had also number of benefits by using various recipes like leaves were being used as a vegetable. The roasted root was used as a coffee adulterant. The root and the leaves were used as appetizer, depurative, digestive, diuretic, laxative and tonic. A decoction of the freshly harvested plant was used for treating gravel; the latex in the stems was applied to warts in order to destroy them. Blood glucose can also be controlled by using powder of dried roots of *Cichorium intybus* L twice in a day. Likewise 4 to 5 leaves of *Zizypus jujube* Mill., were suggested to be chewed daily may help to control blood glucose level. Moreover, powder of *Melia azedarach* L. was proved effective against diabetes if a teaspoon of it used daily before breakfast, for a month.

Further, latex of *Ficus bengalensis* L. leaves and branches mixed with honey and used orally can be effective against diabetes. Another recipe of *Justicia adhatoda* L., was use of its leaves, contain alkaloid, vasicine which has strong bronchodilator activity and used in the form of juice, or decoction to treat respiratory infections. Leaves have antibacterial, anti-inflammatory properties and used externally in the form of poultice for treating wounds and joint pain. *Justicia adhatoda* in combination with *Hyssopus officinalis*, *Sisymbrium irio* Zinn, *Zizypus vulgari*, *Ocimum basilicum*, and *Ephedra vulgaris* are used to make syrup known as ‘Suduri’. Suduri is effective in case of whopping cough, asthma, cold, catarrh, and post influenza cough. Sudari is also a branded product as it is available in market. It promotes expectoration and helps relieving bronchial spasms and brings respiratory functions at normal. Menthol which is extracted from *Mentha* was used in combination of various medications. It provides freshness to breathing. *Morus nigra* L. extract was being used as syrup ‘Sharbat tootseha’ which is very popular and effective against throat infection and pain. Even it is useful in increasing throat glands. Ten milliliters, about 2 teaspoons can be used three to four times’ daily produce effective. Fresh aerial parts of the plant *Solanum nigrum* L. are used to cook as vegetable and are recommended to control diabetes. In addition to that it reduces swelling of liver, spleen, intestines and uterus and regulates the functions. Its extract reduces heat and stimulates urine. Forty milliliters about ¼ tea cup can be used for effectiveness. Aqueous extract of *Oxalis corniculata* L. plant was taken as drink for stomach problem and as cooling agent. The extract of the leaves was used for reducing swelling and redness of eyes and to relieve the pain. The paste of the fresh leaves was used to stop bleeding from wound. These recipes have strong potential to first evaluate them on scientific basis and then transferred them to healthcare products.

### Uniqueness/Novelty of the findings and future prospectus

The study area, Head Maralla is a junction of three Rivers (Jammu Tavi, Manawar wala Tavi, Chenab River) that brings highly fertile soil along with flow of water, desirable for various types of vegetation. Most of the vegetation was site specific like collection of unique plants from four different sites were as follows; at site 1 were *Justicia adhatoda* L., *Lantana camara* L., *Calotropis procera* subsp. Hamiltonii (Wight) Ali, *Ricinus communis* L., *Achyranthes aspera* L., at site 2 were *Canabis sativa* L., *Sonchus asper* (L.) Hill, *Silybum marianum* (L.) Gaertn., *Amaranthus viridis* L., *Cichorium intybus* L., whereas at site 3, *Fumaria indica* (Hausskn.), *Solanum xanthocarpum* Schrad. & H. Wendl., *Typha latifolia* L., *Tribulus terrestris* L., *Melia azedarach* L. Unique plants at site 4 were as *Dalbergia sissoo* DC., *Ficus benghalensis* L., *Acacia nilotica* (L.) Delile, *Zizipus nummularia* (Burm.f.) Wight & Arn., and *Acacia modesta* Wall.

Moreover, some species of bryophytes and pteridophytes (ferns) were the uniqueness of the study area *Azolla pinnata* R. Br., *Marsilea quadrifolia* (aquatic fern), *Osmunda regalis* L., *Marchantia polymorpha* L., and *Riccia cavernosa* Hoffm. Additionally, *Spirogyra buchetii* Petit occurred throughout the season while *Chara Schweinitzii* A. Braun., occurred only during autumn. There was essentially no discussion about medicinal importance rather to document species diversity in Punjab and neighboring areas (Azad Kashmir) [96]. But current studies uncovered the economic importance of these plants, like *Azolla pinnata,* the plant is used as a green manure in paddy fields, in particular to add nitrogen and organic matter to the soil and also act as an activator to speed up the compost-making process. Further, it also possesses ability of phyto-remediation of environmental pollutants. Current findings can lead to develop new form of fertilizers or compost making. *Osmunda regalis* used medicinally as for blood and renal diseases purifier. The roots are used for the production of fibre, and whole plant is eaten as food. Head Maralla is the main water reservoir on the Chenab River just across the Indian border which regulates the water for next heads and barrages like Khanki and Qadirabad. It is an unprotected wetland having wetland biodiversity including various plant species, fish diversity and birds. Birds in the winter are migrated from Siberia to central Asian states from one side and Africa to Middle East from the other side. Another novelty of the study is findings that number of wild plants with new uses for curing different ailments and recipes are being reported from current study. Head Maralla is situated near the foothill of the Himalaya Mountains in both Gujrat and Sialkot districts, therefore, its climate and demography make it hub of different type of vegetation, which require conservation of the area and preservation of indigenous traditional knowledge. Another point is, Head Maralla is a poorly developed local picnic resort where people from central Punjab come and enjoy local hospitality.

Unique plants of a particular site were photographed, collected and preserved for future reference in the herbarium. The area has strong potential for preservation and conservation because it is still in the control of Governmental organizations who can direct all stake holders to act and implement necessary measures for sustainable utilization of wild plant resources. Further, they can increase plantation and reforestation at various localities to safeguard from adverse effects of flood. Various profit making tasks can be initiated to engage local manpower rather to exploit the natural resources are hereby suggested.

### Impact of anthropogenic activities on the area

Since last three decades, different environmental issues have been raised that are resulted by anthropogenic activities as well as natural calamities that affect the plant diversity badly [97]. The anthropogenic activities including urbanization and housing schemes affect the plant community structures. New trends cause ecological changes in vegetation. The increase in human population laid heavy demands such as food and shelters, consequently in the shape of degradation and destruction of floral diversity have observed [98]. Resultantly, agricultural land is gradually reduced which is also one of the causes of the green belt shrinkage. These green belts or patches are now used as built-up areas or commercially available land as a part of urbanization [99]. Moreover, different human malpractices such as blockage of sewage system, irregular garbage dumping, addition of agro-chemicals and many other pollution causing agents are adversely effected biodiversity. These factors have notable severe impact on soil environment, ultimately effected the vegetation [100]. The area is observed with high diversity even contained many unique plant species. Importantly, the area is also unprotected wetland and the birds coming from Siberia, West Asia, Karakorum and Himalayas helps in the seed dispersal and a source of addition of new plant species diversity. There is need to declare the area as protected and to conserve biodiversity for future generations. Current observation declares that population increase, expansion of agricultural lands across river and canal banks, over grazing, fish hunting and over exploitation of resources make it difficult to find biodiversity in safe form and size.

In another study anthropogenic activities have strong influence on the vegetation in the western Himalayas like Naran Valley by plant collection, forest cutting, expansion of agricultural land and grazing pressure[101]. Researchers advocated that indigenous knowledge is culturally valued and scientifically important. They emphasized the wise use and conservation of indigenous knowledge of useful plants that may benefit and improve the living standard of poor people. Similarly, one of the studies reported the existence of 21-28% ethnobotanically important plants in Nepal, out of which about 55% of the flora is of medicinal value [102].

### Conservation of local knowledge and cultural drivers

Conservation of biodiversity is helpful not only for mankind services but also for other organisms, too [103]. For conserving species various protected areas have been declared to conserve species and their habitat. Increase in greenhouse effect, species composition and their habitats are constantly becoming threat to reduce in number [104]. It is noted that protected areas play fundamental role in the safeguarding, protection and longstanding of the biological diversity [105]. Useful plant species required various cultural practices and agronomical requirement for their cultivation, which are difficult to accomplish by one or few institutes only [106]. Further, many of the plants are playing role as cultural drivers and they are used in the form of food, medicine, fodder, timber wood, fuel, cosmetics, spiritual and veterinary purposes. Among 400, 000 total plant species present on the earth, more than 7000 species of higher plants exists in Pakistan, are vascular plants including about 2000 medicinal plant species indicated the potential for benefits concerning next generations [107, 108]. The area is ecologically situated in the Irano-Turanian Region and designated as Western Himalayan Province by Takhtajan [109]. Additionally, the Pakistan possesses a diversity of biotic communities and a relatively rich flora due to great variation in elevation, temperature, precipitation, and other physical parameters [110]. Various studies of ethnobotanical knowledge of Pakistani medicinal plants also supported that old age people showed more folk love for herbal medicines as compared to young ones. The plants were mainly used for stomach, anti-allergic, anti-neuralgia, vermifuge, narcotic and laxative purposes etc., [111].

Study of the 31 trees, herbs and shrubs belonging to 21 families of Chitral Gol National Park (CGNP) Pakistan, revealed the uses as fuel, fence, and as medicine by the local inhabitants. Prominent one are including *Artimisia maritima*, *Artemisia brevifolia*, and *Rosa webbiana* as dominant species suitable for harvesting, while *Ephedra gerardiana* and *Ferula narthex* are also useful [112].

The local government should take steps to stop illegal collection and trade of such valued plants and their seeds. The area is situated in between two districts where a research station and a small scale production unit may be installed to get maximum utilization of the resources, that would be used as revenue generation, along this, postharvest management lacking might be required to safe guard the herbal products and value addition for timber and lumber products.

### Conservation issues of wild medicinal and industrial plants

Due to varied climatic conditions, Head Maralla area is very diverse in biological diversity. More than 150 species are reported (unpublished data) and fauna of different nature including fishes, birds, mammals, amphibians are present in diverse range. Traditional ecological knowledge can play pivotal role in the formulation and implementation of effective conservation strategies at Head Maralla. It can make bridge between natural environment and anthropology which can reflect a long time experience of the wild vegetation of the area and its applications. Currently, few hectares of the land area and rivers are under protection by Government while majority of the area is without any supervision. Therefore, it lacks any conservation strategy to protect wild plants of immense benefits including traditional knowledge. Although various environmental threats such as overgrazing, overexploitations, urbanization asserting negative impact but the exploitation of natural resources without any restriction are big challenge in the area. Moreover, some of the natural disasters like flood and soil erosion are key players against wild plants. The conservation efforts for this area would be fish rearing, fish hunting, cooking, fuel wood sale/purchase, reforestation, nursery business and wildlife hunting with permissions. Following measurement on sustainable basis would enhance the utilization of plants and their products on long term and benefit sharing basis like a) fresh plantation of *Typha*, *Acacia* and *Delbergia* spp., etc., b) catchment areas can be used as fish rearing, c) control grazing, d) nurseries for multiplication of unique species of different sites and concrete based support to river and canal banks. Further ecotourism with community mobilization and benefit sharing can be proved more effective to provide awareness and support such type of activities on sustainable basis.

## Conclusions

The findings of this study certainly open new avenues for research and development. Focus on particular flora shall resultantly bring a systematic approach to cultivate and harvest unique ideas and fruitful outcomes. There is dire need to expand the collection, exploration mission and documentation of the particular flora because in this area seasonal streams and nullahs are flowing which support unique plants to grow in different seasons. Rainfall is highest as compared to all over the country. The area is situated at the foothills of Himalaya and has strong potential to unveil the flora. If such efforts might involve local community, university faculty and students, local government, irrigation department, forest department, Tehsil Municipal Administration, and civil servants for new plantation, cultivation, harvesting and post harvesting, this is assumed to broadcast highest benefits. About 450 hectares of land is being in use and protected by local Forest Department at Head Maralla area, where they have established nurseries and planted different trees like *Eucalyptus*, *Sheesham*, *Dharik*, *Jaman*, and *Shairin* etc. Although plantation is in accordance to forestry laws but they should consider the need and requirement of this area and particular type of vegetation because the sites surrounding head Maralla are hub of unique plant diversity and, fauna and avian fauna. Cultivation of different type of plant species is therefore suggested. Subject area should be declared as protected wetland under Convention on Wetlands Conservation. Topography and climate is also unique that should be owned and efforts to conserve wild flora should also be encouraged. Focused and concentrated efforts are required, which cannot accomplish without the involvement of young literate people. Plant biologist and forest department have ample opportunities to work together and expand plantation across both districts and rivers by motivation and support to enhance fish rearing, control fish hunting, cooking, fuel wood sale/purchase, reforestation, nursery business and wildlife maintenance. In conclusion this document would be a scientific contribution towards preserving traditional knowledge of wild plants of the area and foundation for future conservation efforts.

## Declarations

### Ethics approval and consent to participate

Prior consent of participation was taken (supplementary file attached)

### Consent of publication

NA

### Availability of data and materials

All data generated and/or analyzed during this study are included in this article and its supplementary information files.

### Authors’ contribution

Conceptualization (M.S.I., M.A., M.I.; Analysis M.A.A., S.A.H., N.A., S.M.; Methodology M.A.A., H.M., S.Z.; Resources T.A., N.S., R.M., H.S., Software M.Z., F.B., Data validation K.S., H.A.; MS Drafting & Writing M.S.I., N.F., F.N., Data interpretation A.J.H., S.C., A.W. All authors are agreed to submit it in Plos One for publication.

## Acknowledgements

The cooperation of the local community (elders) is highly acknowledged. Special thanks are extended to army personnel, staff of irrigation and forest department, Number Dar, Chairman Union Council, District Sialkot and Gujrat. Authors are thankful to Prof. Dr. Mushahid Anwar, Chairperson, Department of Geography, University of Gujrat, Gujrat, Pakistan for Head Maralla map cartography. Authors are also grateful to Dr. Bhazad Anwar, Department of English, University of Birmingham, UK, for language editing & proof reading.

## Abbreviations

TRM: Traditional medicines
PHC: Public Health Committee

